# Human NLRC4 can act as a direct sensor for cytosolic flagellin

**DOI:** 10.64898/2026.08.25.746936

**Authors:** Gaopeng Li, Kenyatta Doumanas, Xiao Liu, Nadya Panagides, Liudmila Andreeva, Florian Ingo Schmidt, Clare Elizabeth Bryant, Alexander N. Weber

## Abstract

Innate immune cells sense pathogenic bacteria like *Legionella pneumophila* through patterns such as the protein flagellin, a critical component of the bacterial motility apparatus. Recognition of cytosolic flagellin in mouse immune cells is well understood and mediated by the receptors, neuronal apoptosis inhibitory protein (Naip) 5 or Naip6, which activate the Nlrc4 inflammasome multi-protein complex for initiating cell death or interleukin-1 family cytokine release. However, the role of human NAIP as a cytosolic flagellin sensor remains controversial. Using a multipronged approach, we demonstrate that in a reconstituted cell system human NLRC4 engaged *Legionella* FlaA flagellin directly (i.e. without the need for hNAIP), whereas human NAIP did not interact with FlaA. Ectopic cytosolic FlaA expression also induced NLRC4 oligomerization, a prerequisite for inflammasome activation, in the absence of NAIP. Unexpectedly, the presence of NAIP diminished the binding of NLRC4 to flagellins and subsequent interleukin-1β release. Interestingly, in resting THP-1 cells, NAIP stably interacted with NLRC4, and during infection or stimulation with FlaA pro-inflammatory responses in THP-1 cells were predominantly NLRC4-dependent. Our data highlight NLRC4 as a putative direct sensor of cytosolic flagellins in the human system and NAIP as a potential negative regulator of flagellin sensing.

## Introduction

Immune cells respond to microbial infections by recognizing microbe-associated molecular patterns (MAMPs) through specific receptors, so called pattern recognition receptors (PRRs) ^(Li and Wu 2021).^ MAMPs such as lipopolysaccharide (LPS), a cell wall structural component, and flagellin, a component of the bacterial motility apparatus, can be found in many different types of bacteria, even non-pathogenic commensals, but differ in their PRR activating potential ^(Clasen, Bell et al. 2023)^. Pathogens may additionally contain bona fide pathogen-associated molecular patterns (PAMPs) associated with virulence (e.g. pore-forming toxins and type 3 secretion system, T3SS, components). PRRs are classified in distinct subgroups based on their ligand specificity, function and localization ^(Barnett, Li et al. 2023)^. Nod-like receptors (NLRs) are cytosolic PRRs participating in the formation of inflammasomes – large multi-protein complexes that play a crucial role in triggering the pro-inflammatory response of immune cells. Generally, an inflammasome consists of a sensor protein, which, upon detection of an intracellular MAMP, triggers oligomerization, enabling the recruitment and activation of the effector protein, caspase-1, at times involving further adaptor molecules. The activation of caspase-1 within the now assembled inflammasome complex ultimately results in the maturation and release of the proinflammatory cytokines IL-1β and IL-18 and the formation of membrane pores via cleaved gasdermin (GSDM) proteins, which can lead to cell lysis and further amplification of the inflammatory signal. Among the five inflammasomes described so far, Human NACHT, LRR, and PYD domain–containing protein 3 (NLRP3) is the most prominent, generally monitoring cellular homeostasis by way of detecting efflux of potassium ions via a compromised cell wall (Weber, McManus et al. 2025, Tapia-Abellán, Funk et al. 2026). On the other hand, the neuronal apoptosis inhibitory protein (NAIP)/NLR family CARD domain-containing protein 4 (NLRC4, also known as Ipaf) pathway has the ability to specifically detect and initiate inflammasome activity in response to the MAMP, flagellin, or to PAMP components of bacterial T3SS within the cytosol ^(Kofoed and Vance 2011, Zhao, Yang et al. 2011)^. Both NAIP and NLRC4 contain a central NACHT domain – consisting of nucleotide-binding domain (NBD), helical domain 1 (HD1), winged helix domain (WHD) and helical domain 2 (HD2) – and a C-terminal leucine-rich repeat (LRR) domain. But in their N-termini NAIPs contain baculovirus inhibitor-of-apoptosis repeat (BIR) domains, whereas NLRC4 contains a caspase activation and recruitment domain (CARD), enabling direct engagement of caspase-1 without strict requirement of the adaptor ASC. On the other hand, ASC incorporation may enhance complex stability and amplify cytokine output. (Broz, von Moltke et al. 2010).

Interestingly, mice possess several Naip genes/proteins with different ligand specificities: For example, murine Naip5 was shown to recognize intracellular flagellin and trigger the oligomerization and activation of the murine Nlrc4 inflammasome ^(Kofoed and Vance 2011, Zhao, Yang et al. 2011)^. Consistently, deletion of *Naip5* prevented activation of the murine Nlrc4 inflammasome by flagellin but not by bacterial T3SS components. Subsequently, Naip6, a close homolog of Naip5, demonstrated biochemical and functional characteristics identical to those of Naip5, including the ability to recognize flagellin ^(Lightfield, Persson et al. 2011, Zhao, Yang et al. 2011)^. Instead, activation of the Nlrc4 inflammasome by T3SS components in mice was shown to depend on murine Naip1 and Naip2 ^(Kofoed and Vance 2011, Zhao, Yang et al. 2011, Yang, Zhang et al. 2014, Zhang, Chen et al. 2015)^. For example, murine Naip1 interacts directly with the T3SS needle protein MxiH from *Shigella flexneri ^(Yang, Zhao et al. 2013, Rauch, Tenthorey et al. 2016)^*, whereas murine Naip2 senses T3SS rod proteins such as PrgJ from *Salmonella typhimurium* and BsaL from *Burkholderia pseudomallei* ^(Zhao, Yang et al. 2011)^. Recent structural studies, the ligand engagement of these Naip proteins triggers a conformational change that, in turn, prompts structural rearrangements in Nlrc4 upon its binding to a single ligand-engaged Naip molecule. Nlrc4 is bound in a conformation that enables consecutive assembly with further Nlrc4 molecules into a macromolecular disk that serves as a docking platform for caspase-1 ^(Tenthorey, Kofoed et al. 2014, Zhang, Chen et al. 2015, Paidimuddala, Cao et al. 2023)^. Although a complete structure containing both the bacterial ligand, a Naip molecule and multiple Nlrc4 molecules has not been solved yet, the activation mechanism for murine Naips appears clearly structured around ligand specificity conferred by Naips and inflammasome assembly and execution by Nlrc4 ^(Zhang, Chen et al. 2015, Paidimuddala, Cao et al. 2023)^, i.e. Nlrc4 fulfils a purely structural, but non-sensing role. Activation of the NAIP-NLRC4 inflammasome in mice thus contributes to the expulsion of infected cells from the intestinal epithelium ^(Rauch, Deets et al. 2017)^. Moreover, animals with *Nlrc4* deficiency have been shown to be more susceptible to *S. typhimurium* infections compared to control groups ^(Sellin, Muller et al. 2014)^. Furthermore, *Nlrc4* knockout studies in mice showed increased susceptibility to *Citrobacter rodentium*, thus demonstrating the critical role of the NLRC4 inflammasome host defence strategies in mice ^(Liu, Zaki et al. 2012)^, downstream of Naip sensor proteins.

In contrast to mice, humans, cats, cows and pigs have only a single *NAIP* gene (which is not expressed in pigs) and intact *NAIP* genes are completely absent in dogs, germine and Chinese pangolin ^(Sakuma, Toki et al. 2017, Salova, Sipos et al. 2022)^, indicating that the ‘multi-NAIP’ scenario observed in mice is unique and that other immune systems need to rely on one or no functional NAIP. In the human system, while the intracellular bacterium *Salmonella* Typhimurium was shown to activate the NLRC4 inflammasome in U937 cells, introduction of isolated flagellin or a T3SS rod protein into the cytosol of U937 and THP-1 cells did not ^(Zhao, Yang et al. 2011)^. Subsequently, Feng and colleagues discovered that the needle protein, CprI, but no other components of the T3SS could induce the activation of the human NLRC4 (hNLRC4) inflammasome by binding to hNAIP. Needle proteins from other bacterial species, including *S. typhimurium* and *Shigella flexneri*, were also shown to activate the hNLRC4 inflammasome. This was also confirmed using structural and biochemical studies ^(Matico, Yu et al. 2024)^. These results indicated that hNAIP clearly phenocopies the function of murine Naip1 and 2, and that T3SS component ligand specificity is a NAIP feature shared across species. This, of course, begged the question whether and how sensing of cytosolic flagellin has also converged onto hNAIP or onto another sensor?

In humans, the association of *NLRC4* gain of function mutations with autoinflammatory disease like macrophage activation syndrome (MAS) ^(Canna, de Jesus et al. 2014)^ and neonatal-onset enterocolitis ^(Romberg, Al Moussawi et al. 2014)^ clearly document the physiological relevance of the hNLRC4 inflammasome in patients; but whether and how the hNAIP/hNLRC4 inflammasome is involved in cytosolic flagellin sensing in human cells has been less clear. Generally, biochemical and molecular evidence for the precise detection pathway of cytosolic flagellin in human cells has been far less robust than for the murine system. Previous work indicated that, compared to T3SS components (e.g. PrgI needle protein), hNAIP could not sense flagellin ^(Zhao, Yang et al. 2011, Rayamajhi, Zak et al. 2013, Yang, Zhao et al. 2013)^. Nevertheless, Kortmann et al ^(Kortmann, Brubaker et al. 2015)^ suggested a role of hNAIP in sensing cytosolic flagellin, albeit by only a longer and, until then, overlooked splice isoform of hNAIP, long hNAIP (LhNAIP, cDNA clone NM_004536.2). In their work, *S. Typhimurium*-induced caspase-1 activation, IL-1β release, and cell death were flagellin– and hNAIP/NLRC4-dependent in primary human macrophages which express LhNAIP, whereas U937 and THP-1 macrophage-like cells primarily expressed the short hNAIP (ShNAIP, cDNA clone MGC:177292, IMAGE: 9055275). However, direct interactions of LhNAIP or ShNAIP with flagellin were not shown. Other researchers demonstrated that the response of primary human macrophages to *S. Typhimurium* infection depends on T3SS but not flagellin, using various mutant bacterial strains ^(Reyes Ruiz, Ramirez et al. 2017)^. Collectively, these studies unequivocally confirmed that hNAIP has the ability to bind to cytosolic needle proteins from different bacteria for the activation of the NLRC4 inflammasome. The issue has been complicated further by the fact that under certain circumstances, the NLRC4 inflammasome can functionally interact with the NLRP3 inflammasome ^(Man, Hopkins et al. 2014, Gram, Wright et al. 2021)^, so that cytosolic flagellin sensing in the human system remains somewhat enigmatic.

To address this issue, we here investigated the response of the human NAIP/NLRC4 inflammasome in a simplified heterologous genetic system, namely, HEK293T cells, for the expression of NAIP, NLRC4, T3SS components and different flagellins (e.g. *Legionella* FlaA), in order to avoid possible confounding effects by other inflammasome pathways, e.g. NLRP3, or by bacterial ligand contamination or other co-existing MAMPs. We further validated our results in wild type (WT) and knock out (KO) THP-1 macrophage-like cells. Our findings show that hNLRC4 but not hNAIP – neither LhNAIP nor ShNAIP – interacted with flagellin. Moreover, hNLRC4 oligomerized in the presence of FlaA independently of hNAIP, and hNLRC4 alone was sufficient to trigger in IL1-β and IL-18 release in response to FlaA. Interestingly, increasing amounts of hNAIP reduced FlaA-hNLRC4-mediated IL-1β output, and hNAIP and hNLRC4 constitutively interacted in THP-1 cells, potentially explaining that flagellin responsiveness is constitutively restricted by hNAIP in human cells. Our results thus suggest that hNLRC4 may directly sense the bacterial MAMP, flagellin, but this ability is restricted by hNAIP to favor innate immune responses to T3SS component PAMPs rather than the MAMP flagellin.

## Results

### hNLRC4, but not hNAIP isoforms, can interact with bacterial flagellin in a human cell system

To determine the role of human NAIP (hNAIP) as a cytosolic flagellin sensor, we chose a simple, easy-to-manipulate human cell system devoid of inflammasome proteins, e.g. HEK293T cells, in which inflammasomes can be fully reconstituted by genetic complementation (Kofoed and Vance 2011, Zhao, Yang et al. 2011). Indeed, RT-qPCR and immunoblot analysis of both hNAIP and hNLRC4 confirmed in HEK293T cells exceedingly low mRNA levels for both *hNAIP* isoforms and *hNLRC4* as well as non-detectable protein levels for hNAIP and hNLRC4 (Fig. S1). Thus, the system was ideal to investigate the relative requirements of hNAIP and hNLRC4 with regards to flagellin, which could conveniently be introduced genetically (by ectopic expression) without the risk of confounding by other MAMPs or by other features of an infectious process.

We first explored the possibility of binding between hNAIP and various flagellins from both commensal and pathogenic bacterial species by co-immunoprecipitations (co-IP) in lysates from transiently transfected cells. Flag-tagged, short and long transcripts of human *NAIP* (Flag-ShNAIP and Flag-LhNAIP, respectively), and Flag-tagged mNaip5 (Flag-mNaip5, positive control) were individually co-expressed with HA-tagged flagellin from the commensal bacteria *Bacillus subtilis* (HA-B.SFlic). Immunoblot (IB) analysis of the whole cell lysates (WCL) showed that both NAIPs and flagellin were expressed effectively at the expected molecular weight in all experimental groups. However, despite the fact that HA-B.SFlic was successfully immunoprecipitated (Fig. 1A), neither murine mNaip5, nor any form of human NAIP were pulled down. However, when a plasmid encoding for the HA-tagged flagellin of the pathogenic bacterium *S. typhimurium* (HA-SalFlic) was used (Fig. 1B), Flag-mNaip5 was easily detectable in the pull-down (lane 7), in good agreement with other published data ^(Kofoed and Vance 2011, Zhao, Yang et al. 2011)^. However, Flag-ShNAIP or Flag-LhNAIP could not be detected in the co-transfection groups (lanes 5 and 6). Similar results were obtained for HA-tagged *Legionella pneumophila* FlaA (Fig. 1C): Again, mNaip5 was detected in the IP of the co-transfected group (lane 7), but Flag-ShNAIP or Flag-LhNAIP were not detectable (lanes 5 and 6). These findings suggested that, contrary to expectations and unlike mNaip5, neither LhNAIP nor ShNAIP bound to pathogenic or commensal flagellins in this genetic complementation system.

**Figure 1:**
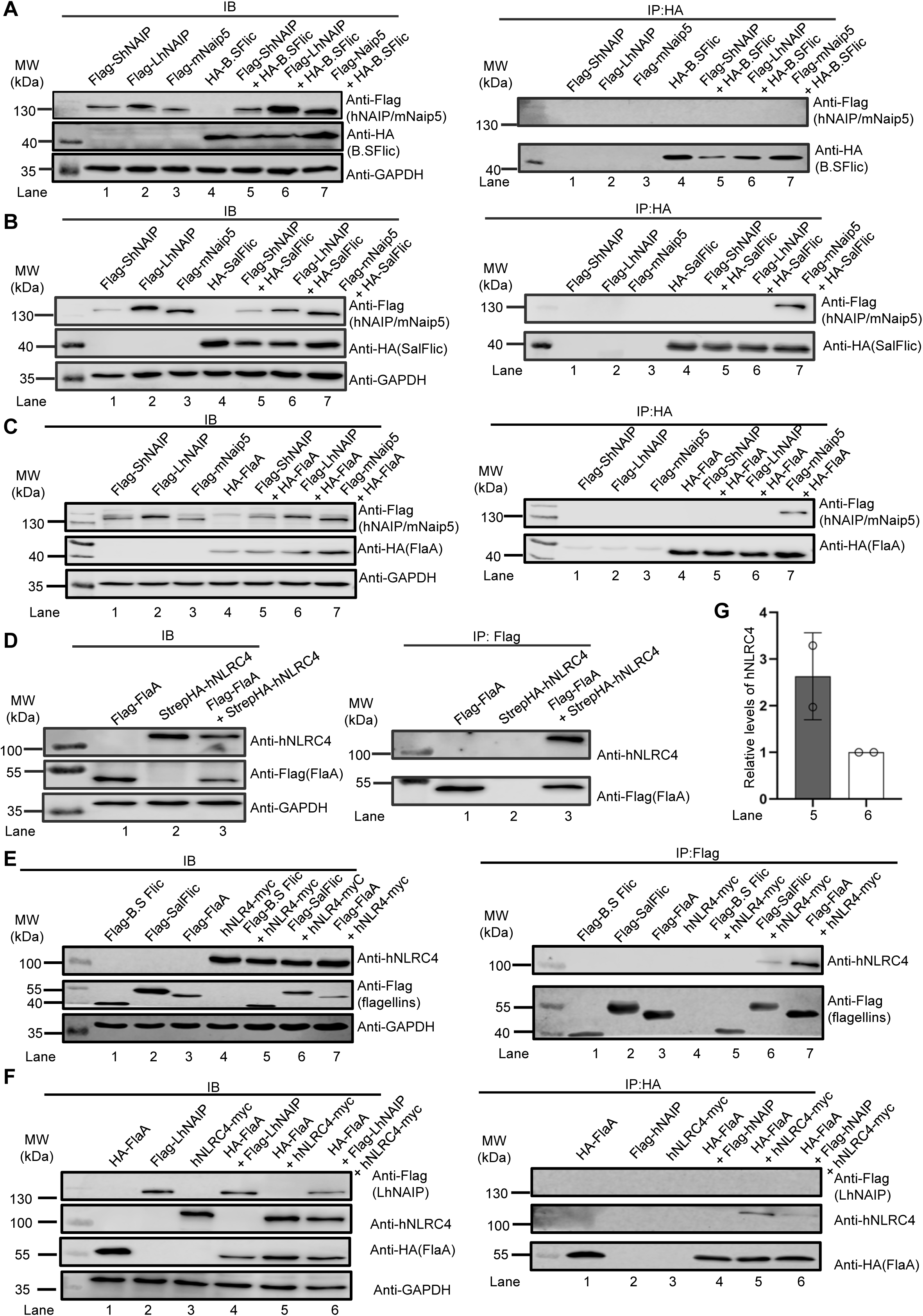
Human NLRC4 but not NAIP binds flagellins from pathogenic bacteria. (**A-C**) Flag-ShNAIP (short human NAIP isoform), Flag-LhNAIP (long human NAIP isoform), and Flag-mNaip5 expression constructs were co-transfected with HA-tagged **(A)** B.SFlic (*Bacillus subtilis* FliC), **(B)** HA-SalFlic (*Salmonella Typhimurium* FliC) or **(C)** HA-FlaA (*Legionella pneumophila* FlaA) expression constructs in HEK293T cells. The HA-flagellins were then immunoprecipitated using anti-HA antibody-conjugated magnetic beads and the presence of NAIPs analyzed by SDS-PAGE and immunoblot. **(D)** Flag-FlaA and StrepHA-hNLRC4 expression constructs were transfected into HEK293T cells as indicated. Immunoprecipitation (IP) was performed with anti-Flag antibody conjugated magnetic beads and both IP and IB fractions were analyzed by SDS-PAGE and immunoblot using the indicated antibodies. **(E)** Flag-tagged flagellins and hNLRC4-myc expression constructs were co-transfected into HEK293T cells. Flag-flagellins were immunoprecipitated with anti-Flag antibody conjugated magnetic beads and hNLRC4 detected in the co-transfection groups by SDS-PAGE and immunoblot using the indicated antibodies. **(F)** HA-FlaA, Flag-hNAIP, and hNLRC4-myc expression constructs were co-transfected into HEK293T cells. Flag-flagellins were immunoprecipitated with anti-Flag antibody conjugated magnetic beads and hNLRC4 and hNAIP detected in the co-transfection groups by SDS-PAGE and immunoblot using the indicated antibodies detected. **(G)** Quantification of the intensity of hNLRC4 bands in lanes 5 and 6 of the IP group shown in (F). In A-C, F (n=2), D, E (n=3) one out of n representative experiments is shown; G (n=2) shows combined data from n experiments.

Since we could not detect binding between hNAIP and flagellins but flagellin responses in human cells had been suggested to nevertheless be NLRC4-dependent ^(Kortmann, Brubaker et al. 2015)^, we investigated whether human NLRC4 itself might have the ability to interact with flagellins. Flag-tagged FlaA was therefore co-transfected with StrepHA-tagged hNLRC4 (Fig. 1D). Conversely to hNAIP, and somewhat surprisingly, hNLRC4 was co-immunoprecipitated with FlaA (lane 3), indicating that hNLRC4 might have the capability to bind to FlaA in the absence of hNAIP. Next, we tested other flagellins, namely Flag-tagged B.SFlic, Flag-tagged SalFlic, and Flag-tagged FlaA in parallel with a myc-tagged hNLRC4 expression construct, respectively (Fig. 1E). Interestingly, hNLRC4 was co-immunoprecipitated with all flagellins except for Flag-B.SFlic (lanes 5,6 and 7), suggesting that similar to mNaip5, hNLRC4 binds to flagellins from pathogenic bacteria but not commensal bacteria. Interestingly, in the IP samples, the intensity of the band of hNLRC4 was stronger in the *Legionella* FlaA co-transfection group compared to the *Salmonella* FliC co-transfection group, indicating that potentially the binding affinity between hNLRC4 and *Legionella* flagellin might be higher than the affinity between hNLRC4 and *Salmonella* flagellin. Since hNAIP failed to bind flagellins in our system, we sought to investigate whether it could influence the binding of cytosolic flagellin to hNLRC4. Consequently, HA-FlaA, Flag-LhNAIP, and hNLRC4-myc constructs were transfected together into HEK293T cells (Fig. 1F). Although there was still an interaction of hNLRC4 and FlaA in the presence of hNAIP, the detected amount of pulled down hNLRC4 in the triple transfection group (lane 6) was weaker than that in the absence of hNAIP transfection (lane 5, quantified in Fig. 1G). In summary, these data suggested that, similar to mNaip5, hNLRC4 can interacted with flagellins from pathogenic bacteria in the absence of NAIP which appeared to not interact with any flagellins tested and rather decreased the interaction between flagellins and hNLRC4.

### hNLRC4 alone is sufficient for FlaA-induced reconstituted inflammasome assembly

Previous data showed that, after binding to their specific ligands, murine Naips trigger the formation of mNlrc4 oligomers in the HEK293T reconstituted system ^(Kofoed and Vance 2011, Zhao, Yang et al. 2011)^. Therefore, we wondered whether, in the absence of NAIP, flagellins could trigger the oligomerization of hNLRC4, a hallmark of NLRC4 inflammasome activation ^(Yang, Zhang et al. 2014)^. A HEK293T-based, fully reconstituted inflammasome system, involving transfection of not only trigger (HA-FlaA) and NLRs (Flag-LhNAIP, myc-hNLRC4) but also pro-hCaspase-1 and pro-hIL-1β system, was previously described and combined with native PAGE, a method able to identify e.g. oligomerized murine Nlrc4 as “smears” visible upon sensing of cytosolic flagellins and other T3SS components ^(Kofoed and Vance 2011, Yang, Zhao et al. 2013)^. Applying this to human NLRC4 (Fig. 2A), in the absence of FlaA transfection, no band of hNLRC4 with high molecular weight was detectable (lane 1). However, in the presence of FlaA, but even without hNAIP transfection, oligomers of hNLRC4 could be detected (lane 2). This was specific for hNLRC4 as a negative control condition lacking hNLRC4 transfection but containing hNAIP did not show a smear (lane 3). Compared to lane 2, the intensity of the smear band of hNLRC4 oligomers was reduced in lanes 4 and 5, indicating that the oligomerization of hNLRC4 is decreased in the absence of pro-hCaspase-1 and pro-hIL-1β. Consistently with the co-IP result (*cf.* Figs. 1F and 1G lane 5 vs 6), the intensity of the hNLRC4 oligomer smear was also decreased in lane 6 when NAIP was transfected in addition to all other NLRC4 inflammasome components. To check whether HA-FlaA was incorporated into the hNLRC4 oligomer complex, we also used an anti-HA antibody to detect a possible shift of FlaA to a larger molecular weight “smear” (Fig. 2B). No shift was observed in lane 1, which served as the negative control, as the transfection mix did not contain FlaA. Interestingly, a FlaA “smear” was observed in lane 2, i.e. in the presence of FlaA and hNLRC4 but in the absence of hNAIP transfection. However, the shift was no longer observable in lane 3 (without hNLRC4 transfection) and thus specifically dependent on hNLRC4. In lanes 4, 5, and 6, which all contained hNLRC4, a recurrence of the smear was visible, indicative of a shifting of FlaA, which was considerably more pronounced in lanes 4-6 (with LhNAIP) compared to lane 2 (no LhNAIP). Finally, the experiment was repeated using anti-NAIP antibodies (Fig. 2C), to ascertain whether hNAIP was recruited to a presumed FlaA– and hNLRC4-containing complex. However, the anti-NAIP antibodies, despite being specific (*cf.* Fig. S1D), did not highlight a smear in any of the conditions. These observations suggest that FlaA directly interacts with a hNLRC4 oligomeric complex, and that this process is independent of hNAIP. Conventional SDS-PAGE and IB analysis confirmed the correct expression of all constructs across all the experimental groups (Fig. 2D). In an orthogonal approach, we also attempted to visualize oligomers of hNLRC4 as a consequence of FlaA binding by confocal fluorescence microscopy and quantification. FlaA and hNLRC4 were expressed as fusions with mCherry and eGFP in distinct transfection groups (Fig. 2E). As expected, mCherry-FlaA and eGFP-hNLRC4 were uniformly distributed throughout the cytoplasm in HEK293T cells only transfected with one or the other (panels of group 1 and 2 in Fig. 2F). However, puncta indicative of hNLRC4 oligomers ^(Man, Hopkins et al. 2014)^ were observed in both mCherry-FlaA– and eGFP-hNLRC4 co-transfected groups (panel group 3), even in the absence of hNAIP (panel group 4), albeit at lower levels. In the absence of FlaA (panel group 5), there were no puncta of hNLRC4, suggesting that the formation of oligomers of human NLRC4 in this assay was indeed FlaA-dependent. These conclusions were confirmed by quantification of the microscopy data (Fig. 2G): The proportion of cells exhibiting hNLRC4 oligomerization was very low in single transfection group 1. However, it was significantly higher in group 3, when LhNAIP was absent. These microscopy results, which were fully consistent with the native PAGE data, thus suggested that in the presence of FlaA, hNLRC4 can directly form oligomers, a hallmark of inflammasome assembly, even in the absence of NAIP. In fact, the presence of LhNAIP seems to inhibit the oligomerization of hNLRC4.

**Figure 2:**
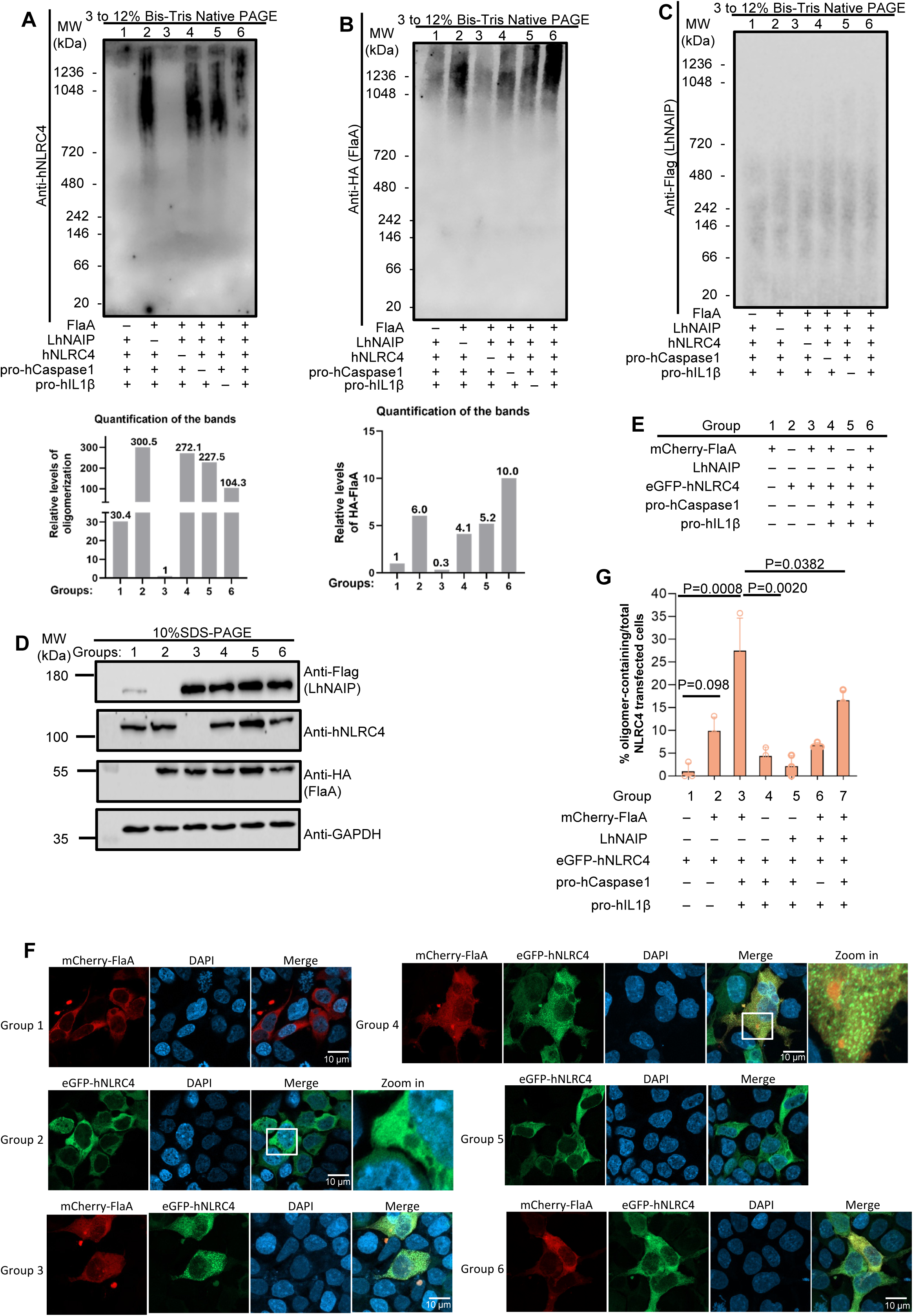
FlaA triggers hNLRC4 oligomerization independently of hNAIP. (**A–C**) hNLRC4 inflammasome system reconstitution in HEK293T cells. HA-FlaA, Flag-LhNAIP, myc-hNLRC4, pro-hCaspase-1 and pro-hIL-1β expression plasmids were co-transfected as indicated, cells lysed under native conditions, and lysates analyzed by native PAGE and immunoblotting using the indicated antibodies, in **(A)** anti-hNLRC4, in **(B)** anti-HA antibody and in **(C)** anti-Flag antibodies. **(D)** Equivalent SDS-PAGE and immunoblot of the same lysates with the indicated antibodies. **(E)** Schematic overview of the group transfection setup for **(F)** Airyscan confocal microscopy images of each group shown in (E). FlaA was tagged with mCherry (red pseudocolor), and hNLRC4 was tagged with eGFP (green). Nuclei were stained using Hoechst (blue). **(G)** Blinded observer quantification of the fraction of cells transfected with eGFP-hNLRC4 containing oligomers of eGFP-hNLRC4 in each tiled image, i.e. the number cells with oligomers was divided by the total number of eGFP-hNLRC4-transfected cells. Each dot one tiled image (mean+SD). In A-F one representative of n=3 experiments is shown, G shows data combined from n=3 (one – way ANOVA).

### FlaA triggers the release of hIL-1β from NLRC4 inflammasome-reconstituted cells in the absence of NAIP

Based on the above findings, it was imperative to examine in our reconstituted system whether flagellin-mediated oligomerization of hNLRC4 could induce the release of hIL-1β, the terminal outcome of hNLRC4 inflammasome activation ^(Kofoed and Vance 2011, Zhao, Yang et al. 2011)^, in the absence of NAIP. As shown in Fig. 3A, HEK293T were transfected with different constructs in various combinations, with group 4 carrying all of them. For hNAIP, we used both the LhNAIP construct (Figs. 3B-D) and the ShNAIP construct (Fig. S2) In addition, different types of flagellins were tested. The levels of hIL-1β were generally quite low but comparable in the three groups in which B.SFlic was transfected (Fig. 3B), consistent with the co-IP results (*cf.* Fig. 1A) suggesting that the expression of B.SFlic may not be able to trigger activation and release of hIL-1β. However, hIL-1β release above background was measurable when SalFlic was transfected (Fig. 3C), even in the absence of LhNAIP. In the presence of LhNAIP (group 4), however, the release of hIL-1β was comparable. The levels of hIL-1β induced by FlaA in the reconstituted system were similar to those of SalFlic, but here the effect of the absence (group 2) or presence (group 4) of LhNAIP in combination with FlaA showed a pronounced effect (Fig. 3D), namely that IL-1β release was lower when hNAIP was co-transfected. Similar results were obtained for ShNAIP (Figs. S2A and S2B). To strengthen these conclusions, the dependence on caspase-1 as a terminal executor of the cascade was also checked (Figs. 3E-H). This confirmed the full dependence of IL-1β release in this system on caspase-1, FlaA and hNLRC4, but not hNAIP, whose presence rather dampened IL-1β release.

**Figure 3:**
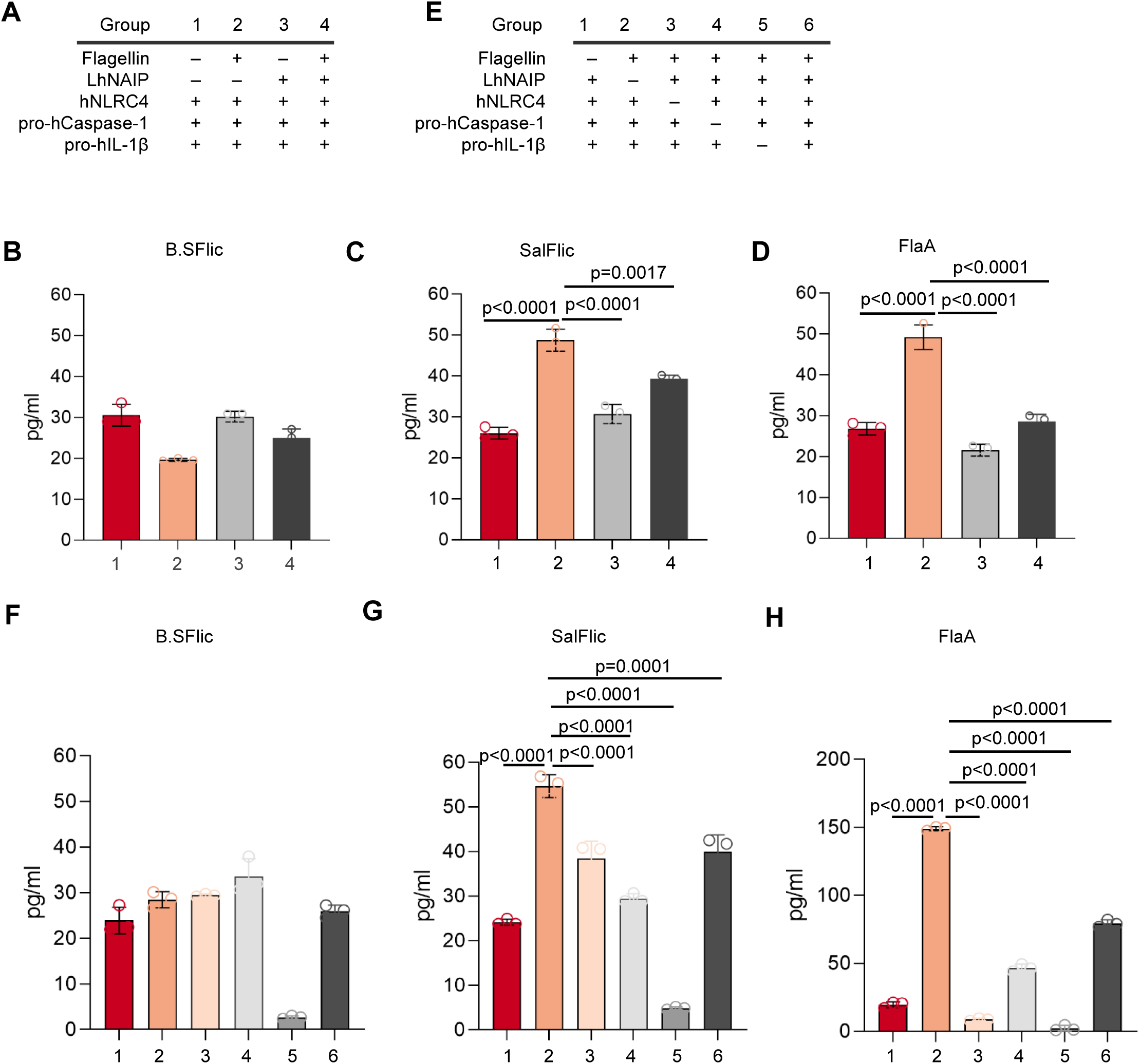
Flagellin-mediated IL-β release in the reconstituted hNLRC4 inflammasome system. (**A-D**) hNLRC4 inflammasome system reconstitution in HEK293T cells. HA-flagellins, Flag-LhNAIP, myc-hNLRC4, pro-hCaspase-1 and pro-hIL-1β expression plasmids were co-transfected as indicated and after 48 h hIL-1β was quantified by triplicate ELISA. **(E-H)** As in AD but with HA-flagellins additional combinations/conditions. In A-H data are representative of n = 3 independent experiments (each dot represents one technical replicate, mean+/-SD plotted, Student’s t-test).

### Human NAIP interacts with human NLRC4 in the absence of flagellin and inhibits the reconstituted hNLRC4 inflammasome

Based on the above findings, the presence of hNAIP seemed to somehow impair the hNLRC4-dependent effects of cytosolic pathogenic flagellins in the reconstituted inflammasome system. The simplest explanation for this observation is that hNAIP has the ability to interact with hNLRC4 directly and block its function. To determine whether hNAIP is capable of binding to hNLRC4, Flag-ShNAIP and Flag-LhNAIP expression constructs were co-transfected with hNLRC4-myc into HEK293T cells for co-IPs (Fig. 4A). Indeed, hNLRC4 was detectable in both the co-transfection IPs, suggesting that both ShNAIP and LhNAIP have the ability to directly bind to hNLRC4 in this overexpression system. To confirm this interaction in a more physiological setting, we turned to THP-1 cells, where both hNAIP and hNLRC4 are endogenously expressed (*cf.* Fig. S1). hNAIP and hNLRC4 knockout THP-1 pools generated by Cas9-CRISPR (see Methods) served as antibody specificity controls (Fig. S3). After pulling down endogenous hNLRC4 using anti-hNLRC4 antibody, endogenous hNAIP was detected in WT THP-1 but not in hNAIP KO THP-1. As expected, hNLRC4 was not detected in hNLRC4 KO cell line lysates after incubation with anti-hNLRC4 antibody, further confirming the specificity of the antibody used for the IP. Neither hNLRC4 nor hNAIP could be found in the isotype control groups (Fig. 4B). Collectively, this data demonstrated that hNAIP has the ability to interact with hNLRC4 directly, even in the absence of flagellin stimulation, at least in transfected HEK293T and endogenously in THP-1 cells. Based on the prior observation that the presence of hNAIP somehow dampens hNLRC4– and flagellin-dependent release of IL-1β, we speculated that hNAIP could block the release of hIL1-β in the reconstituted system. To test this hypothesis, we measured the levels of hIL1-β in the reconstituted system in the presence of increasing concentrations of either the ShNAIP or the LhNAIP construct (Figs. 4C and 4E). We found that the higher ShNAIP expression (confirmed by immunoblot, Fig. 4D), the lower the levels of hIL-1β released in the presence of FlaA (groups 5, 6, and 7), with similar results obtained for LhNAIP (Figs. 4E and F) Of note, increasing expression of another NLR, NLRP6, or the unrelated control protein, GFP, did not have this effect (Figs. 4G, 4H and Fig. S4).These results suggest that the blocking effect observed for ShNAIP and LhNAIP on FlaA-hNLRC4-dependent IL-1β release was relatively specific.

**Figure 4:**
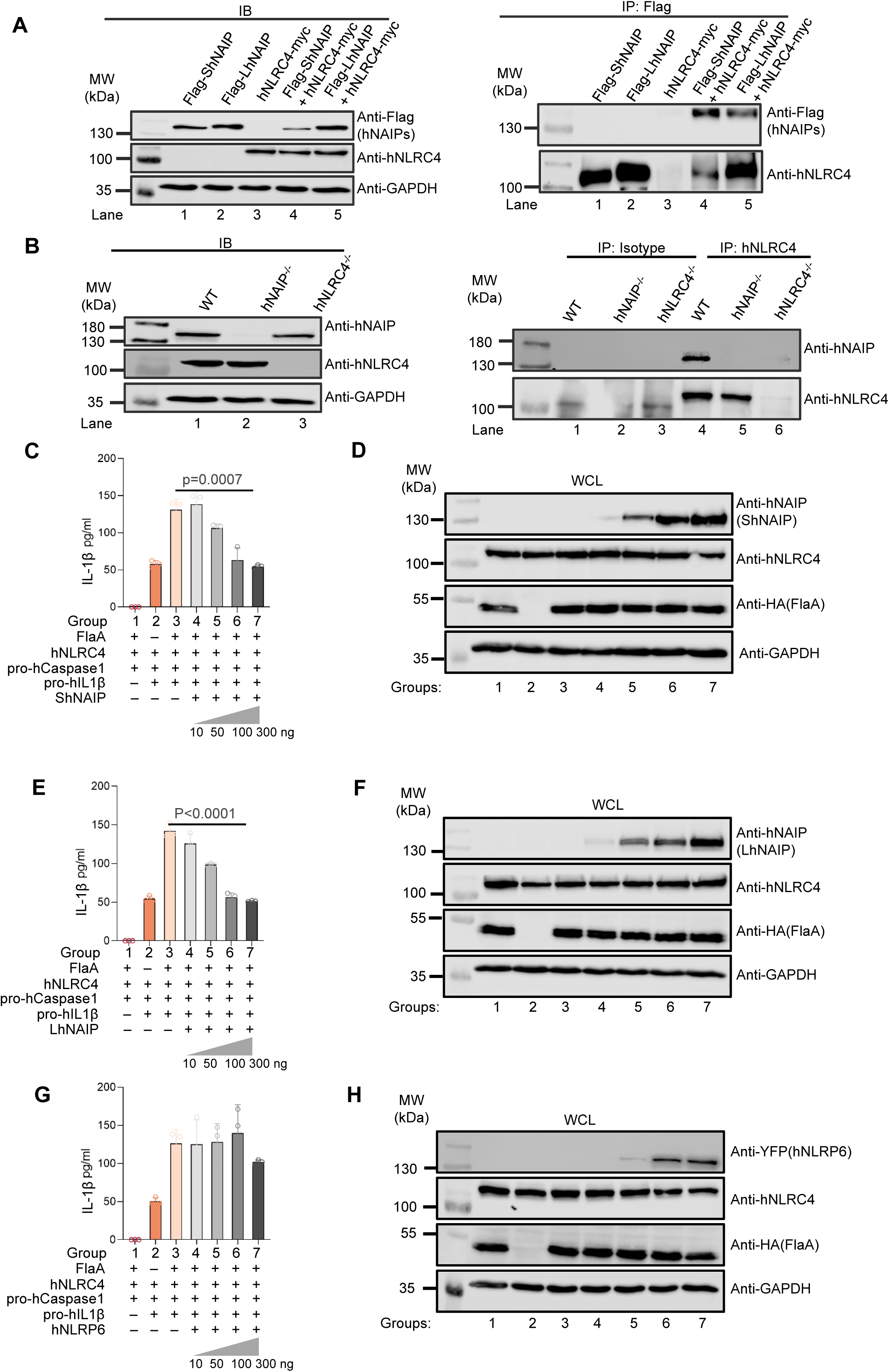
hNLRC4 interacts with and IL-1β release is inhibited by hNAIP. **(A)** Flag-hNAIP and myc-hNLRC4 constructs were co-transfected into HEK293T cells, followed by IP using anti-Flag magnetic beads. Both WCL and IP samples were analyzed using SDS-PAGE and IB using the indicated antibodies. **(B)** The indicated WT and knockout THP-1 cell lines lysed for IP using anti-hNLRC4 antibodies, and WCL and IP samples analyzed by SDS-PAGE and IB to detect endogenous hNAIP using the indicated antibodies. **(C)** as in Fig. 2 but the ShNAIP plasmid amount was gradually increased from group 4 to 7. **(D)** Lysates from C analyzed by SDS-PAGE and IB. **(E)** as in C but with LhNAIP. **(F)** Lysates from E analyzed as in D. **(G)** as in C but with an increased concentration of human NLRP6 plasmid instead of hNAIP plasmid. **(H)** Lysates from G analyzed as in D. In A and B one representative out of n=2 biological replicates is shown. C-H are representative of three experiments, n = 3, mean+SD; each dot represents one technical replicate, Student’s t-test.

### Endogenous hNLRC4 is activated by FlaA and flagellated bacterial pathogens in THP-1 cells

The findings in HEK293T cells so far provided compelling evidence that hNLRC4 functions as a primary receptor for cytosolic flagellins, and hNAIP may rather inhibit this sensing mechanism by binding to hNLRC4 directly. To confirm that a similar mechanism operates in immune cells, we performed further experiments in the aforementioned WT, hNAIP KO and hNLRC4 KO THP-1 cells (*cf.* Fig. S3). For this, we exploited the ability of the protective antigen (PA) and the N-terminal domain of the lethal factor (Lfn) of the lethal toxin of *Bacillus anthracis* to deliver heterologous proteins in the cytosol of mammalian cells ^(Kushner, Zhang et al. 2003)^. As expected, both hNAIP KO and hNLRC4 KO THP-1 cell lines both lost the ability to respond with IL-1β and IL-18 release to cytosolic T3SS needle and MxiH proteins (Figs. S5A and S5B). We then tested their response of WT and KO THP-1 cell lines to cytosolic FlaA conjugated to Lfn alone or in combination with PA, with Lfn-Needle alone or in combination with PA serving as the positive control (Fig. 5A).When compared to the non-treatment group, WT exposed to Lfn-FlaA showed an increase in the levels of IL-1β after 72 h of incubation (Fig. 5B), albeit far less than PA combined with Lfn-Needle or Lfn-MxiH, in line with the reported lower effectiveness of PA delivery for the relatively large (48 kDa) FlaA compared to the small needle protein like *S.* Typhimurium needle PrgI (8.9 kDa) ^(Yang, Zhao et al. 2013)^. In contrast, hNLRC4 KO THP-1 cells released even lower levels of IL-1β levels in response to PA plus Lfn-FlaA compared to WT cells, confirming dependence on hNLRC4. Since hNAIP KO cells showed elevated levels compared to WT cells, we can conclude that FlaA sensing is not hNAIP dependent, but the removal of hNAIP rather augments the presumably still NLRC4-dependent response. Similar results were obtained for the release of IL-18 (Fig. 5C), although the levels of this cytokine were generally lower than IL-1β levels and an increase of hNAIP-deficient over WT cells was not evident. This data is consistent with earlier results indicating that the FlaA response relies exclusively on NLRC4 as sensor and not hNAIP.

**Figure 5:**
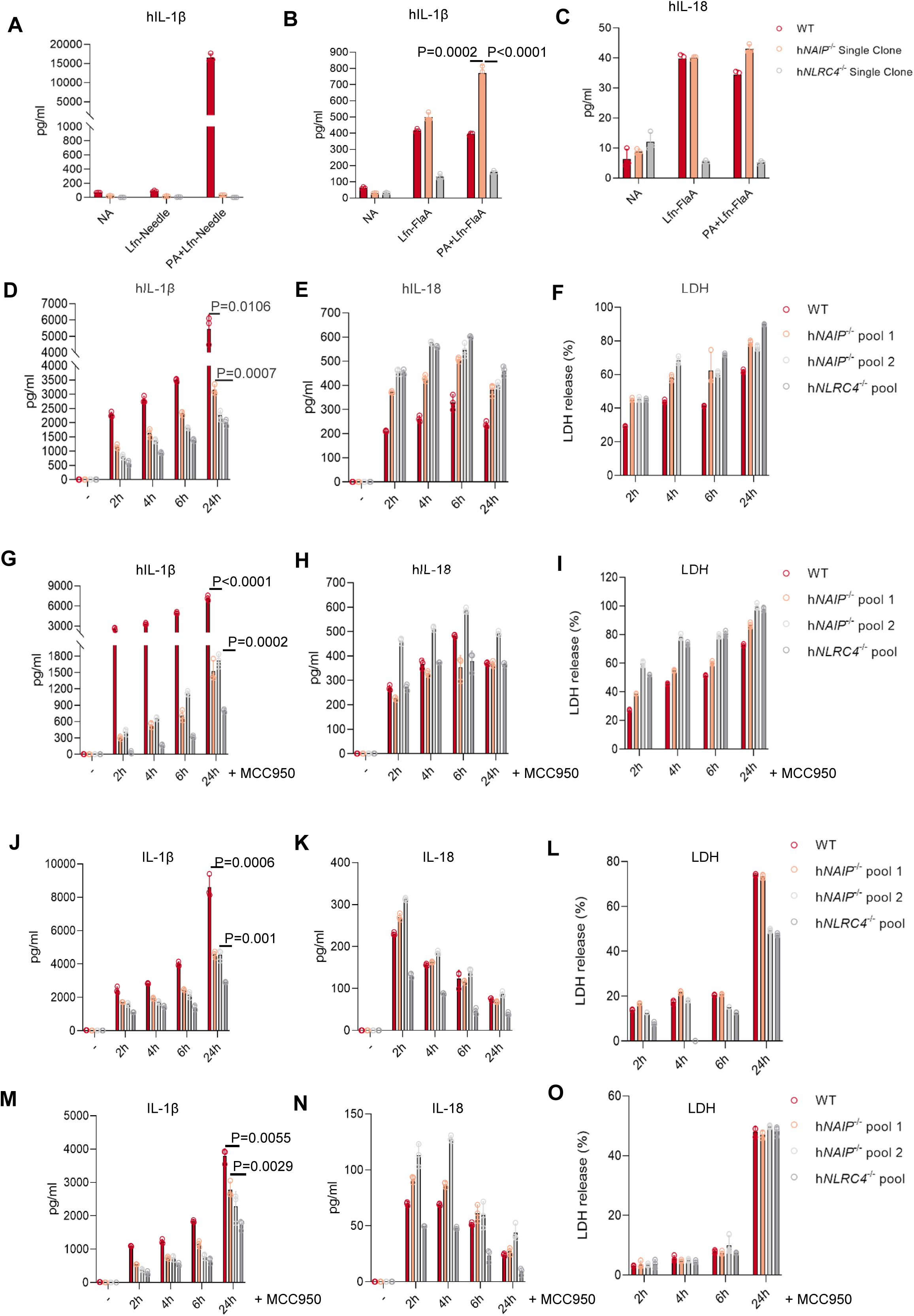
Cytokine release of WT and knockout cell lines in response to FlaA and flagellated bacteria. (**A-B**) WT and KO PMA-differentiated THP-1 cells were treated with either Lfn-FlaA or the combination of PA plus Lfn-FlaA. 72 later, hIL-1β (A) and hIL-18 (B) levels were quantified by triplicate ELISA**. (C-E)** WT and KO PMA-differentiated THP-1 cells were infected with *S. Typhimurium* at MOI 10. At the indicated time points, the concentration of cytokines and LDH was measured in technical triplicates. **(F-H)** As in C-E but with 10 μM MCC950 added 1 h prior to infection. **(I-K)** as in C-E but with *L. pneumophila* at MOI 50. **(L-N)** As in I-K but with 10 μM MCC950 added 1 h prior to infection. In A-K (n=3) are representative of n experiments, each dot represents one technical replicate (mean +/-SD, Student’s t-test).

### hNLRC4 and hNAIP dependence in a complex setting: infection

To further test this model, we investigated the response of WT and KO THP-1 cells to infection with *Salmonella* Typhimurium, bearing in mind that these bacteria contain multiple additional PRRs and both MAMPs like flagellin and PAMPs like PrgI/PrgJ ^(Zhao, Yang et al. 2011, Yang, Zhao et al. 2013, Naseer, Egan et al. 2022)^. According to our hypothesis, hNLRC4 KO deletion would be expected to abrogate both MAMP (flagellin) and PAMP (T3SS component) sensing, whereas hNAIP KO should only affect the latter, whilst potentially augmenting the former. Overall output would thus depend on the relative abundance and affinity of both PRR activators and might also be masked by activation of other, unrelated PRRs in the process. Despite this expected difficulty in the interpretation, cells were infected with either bacterium at MOI 10 (*Salmonella*). At designated time points the culture supernatant was collected, and the released IL-1β, IL-18, and LDH (a proxy for cell death) quantified. In this setting, compared to WT, the deficiency of both hNAIP or hNLRC4 significantly decreased the levels of hIL-1β. However, the level of hIL-1β in hNAIP KO pools was clearly higher than that in hNLRC4 KO pool (Fig. 5D). This supports our previous findings that hNAIP does not recognize flagellin as otherwise, hNAIP and hNLRC4 KO should be indistinguishable here, similar to the results obtained for LFn-Needle or LFn-MxiH stimulation (*cf.* Fig. S5). Rather, the deficiency of hNAIP only partially diminished the cell response in the context whole bacterial infection, which is consistent with published data showing that hNAIP recognizes T3SS needle ^(Yang, Zhao et al. 2013)^. According to our previous results, this would indicate that needle PAMP sensing is dominant over flagellin MAMP sensing. However, hIL-18 (Fig. 5E), which was generally very low compared to IL-1β, was not reduced in the KO lines but rather slightly increased. The LDH pattern was similar to that of hIL-18 (Fig. 5F). As THP-1 *S.* Typhimurium infection was also reported to activate NLRP3 ^(Gram, Wright et al. 2021)^, we speculated that, concomitant NLRP3 activation might contribute to these cellular responses. We therefore repeated the infection in the presence of the NLRP3 inhibitor MCC950 ^(Coll, Robertson et al. 2015)^, and found that IL-1β levels in response to *S.* Typhimurium infection were uniformly reduced in all KO cell lines, but the differences between WT, hNAIP and hNLRC4 clones observed before were still consistent (Fig. 5G), namely that hNAIP KO, albeit being lower then WT, responded more readily than hNLRC4 KO. Under MCC950 inhibition, the released IL-18 levels were increased for hNAIP pool 2 whereas hNAIP1 and hNLRC4 were comparable to WT (Fig. 5H). Unlike IL-1 and IL-18 release, LDH release, and thus cell death, appeared unaffected by MCC950, suggesting that cytokine maturation and cell death in THP-1 cells are not necessarily NLRP3, hNLRC4 or hNAIP dependent. We next compared this to another bacterial pathogen with less PAMPs compared to *Salmonella*, and in which flagellin is considered a dominant MAMP, namely *Legionella pneumophila* ^(Zhao, Yang et al. 2011)^. Consequently, the cells were infected (MOI 50) with *L. pneumophila*, with and without MCC950. As before, levels of IL-1β increased over time but the graded reduction observed before was also apparent, albeit less pronounced: hNAIP KO clones appeared clearly reduced in their ability to releasee IL-1β and hNLRC4 KO cells produced the least IL-1β (Figs. 5J and 5M). If hNAIP was strictly required, we would have expected a similar reduction. Interestingly, IL-18 release (Figs. 5K and 5N) showed a pattern more similar to PA+Lfn-Fla stimulation (*cf.* Fig. 5B/C) in that IL-18 release by the two hNAIP KO pools was higher than in WT cells, whereas in hNLRC4 KO cells IL-18 release was again lower than WT cells. In this infection setting, MCC950 reduced cell death <24 h more than 50% (Figs. 5N vs 5K), suggesting it is predominantly NLRP3 dependent and difficult to relate to hNAIP or hNLRC4. In conclusion, despite the complexity of an infection system in multi-PRR cells using multi-MAMP/-PAMP pathogens, we can conclude that hNAIP loss does not phenocopy hNLRC4 deficiency, which indirectly confirms that hNAIP is not the sole sensor of all murine Naip ligands including flagellin; rather the difference between hNAIP KO and hNLRC4 KO cells, especially for the more flagellin-dominant *Legionella* are consistent with data from the HEK293T system and Lfn-FlaA-only stimulation indicating that hNLRC4 senses flagellin and hNAIP might restrict its activity.

## Discussion

In this study we provide data indicating that in the human system hNLRC4, and not hNAIP, has the ability to bind and trigger responses to certain cytosolic flagellins, e.g. from *Salmonella* and *Legionella*, but not flagellins from commensal bacteria. Thus, hNLRC4 and not hNAIP could be immunoprecipitated by *Salmonella* FliC and *Legionella* FlaA. In the presence of these flagellins, hNLRC4, and not hNAIP, formed oligomeric higher MW complexes and puncta indicative of inflammasome complex assembly. Consistently, release of hIL-1β from inflammasome reconstituted cells was observed in the absence of hNAIP but only when hNLRC4 was present. If anything, when hNAIP was present, high MW complex and puncta formation, as well as IL-1β release was diminished. Given that hNAIP can bind hNLRC4 constitutively, this suggests that hNAIP, whilst clearly acting as a sensor for T3SS components, might be a negative regulator of hNLRC4 activation by flagellin in the human system. Our infection results are less stringent to interpret due to the inherent complexity of timing and the concomitant of multiple MAMPs and DAMPs; nevertheless, especially for *Legionella*, they provide indirect support as a unique and exclusive sensing role for flagellin and T3SS would have stipulated to see no differences between hNAIP KO and hNLRC4 KO as was the case for the bona fide hNAIP-dependent activators, LFn-Needle or LFn-MxiH (*cf.* Fig. S5). Further infection experiments with genetically modified *Salmonella* that express only certain MAMPs or PAMPs might be of interest but since flagellin expression is co-regulated by T3SS (e.g. SPI-1) ^(Naseer, Egan et al. 2022)^ this is difficult to accomplish and outside the scope of this present work.

Of course, these findings beg the question how NLRC4 might directly sense flagellin? Formal proof for a direct interaction is difficult to attain using cellular systems. Thus, future biochemical and structural studies prompted by our results and using recombinant proteins may have to shed light on this issue. However, based on the structural information available to date, a direct interaction seems plausible: The crystal structure of the mNaip5-FlaA complex revealed the N-terminal helix, BIR1, HD1, HD2, ID, and LRR regions of Naip5 to contribute to the formation of a hydrophobic pocket that wraps around the FlaA D0 domain ^(Yang, Zhao et al. 2013, Tenthorey, Kofoed et al. 2014, Paidimuddala, Cao et al. 2023)^. The crystal structure also confirmed that the C-terminal 35 residues of flagellin are both necessary and sufficient for activating mNaip5. Notably, the NBD and WHD, which are essential for the assembly of the mNaip5-Nlrc4 inflammasome, do not contribute to the detection of the FliC D0 by mNaip5. Alignment of the hNAIP and murine Naip protein sequences revealed that hNAIP is closest to mNaip2, i.e. the needle-sensing mNaip (Fig. S6A), in good agreement with the published ^(Zhao, Yang et al. 2011, Rayamajhi, Zak et al. 2013, Yang, Zhao et al. 2013)^ and observed essential requirement of hNAIP for sensing T3SS components in the human system (*cf.* Fig. 5A and S5). Moreover, flagellin-interacting residues of mNaip5 are largely conserved in mNaip6 but not in mNaip2 (not shown), clarifying why mNaip5 and mNaip6 selectively detect flagellin. From a sequence point of view, the similarity of hNAIP to mNaip2 but not mNaip5/6 argues that only T3SS, and not flagellin binding, is conserved in hNAIP. Human NLRC4 also contains NBD, NACHT, WHD and LRR which are conserved among NLRs (Fig. 6A). Aligning the crucial residues in these domains of mNaip5 with amino acids in the corresponding domains of hNLRC4 reveals a relatively high similarity between mNaip5 and human NLRC4 (Figs. 6A, 6B and S6B). For example, residues R846, L848 and W849 in the HD2, and residues F964 and E968 in the ID of mNaip5 are identical to those in the respective domains of hNLRC4. Moreover, I971, K972, N973, Y974, E975, and N976 in the ID and C1329, H1360, and S1363 in LRR of mNaip5 have similar properties to the corresponding residues in hNLRC4. We concede that these results do not prove, but could indicate, that hNLRC4 has the potential to form the same hydrophobic pocket to bind to flagellins as mNaip5 does. Moreover, the assembly mechanism for hNLRC4 upon engagement of *Bacillus thailandensis* needle protein by hNAIP relies on the activated hNAIP providing an extended L736 as a ‘key’ to unlock the closed hNLRC4 conformation ^(Matico, Yu et al. 2024)^. Thereafter, open hNLRC4 is sufficient to open up additional closed hNLRC4s to assembly. It appears conceivable that interaction with FlaA might provide a similar cue and an hNLRC4 ‘opened’ by FlaA binding be sufficient to trigger the assembly of a full active hNLRC4 oligomer. However, this prediction awaits experimental validation.

**Figure 6:**
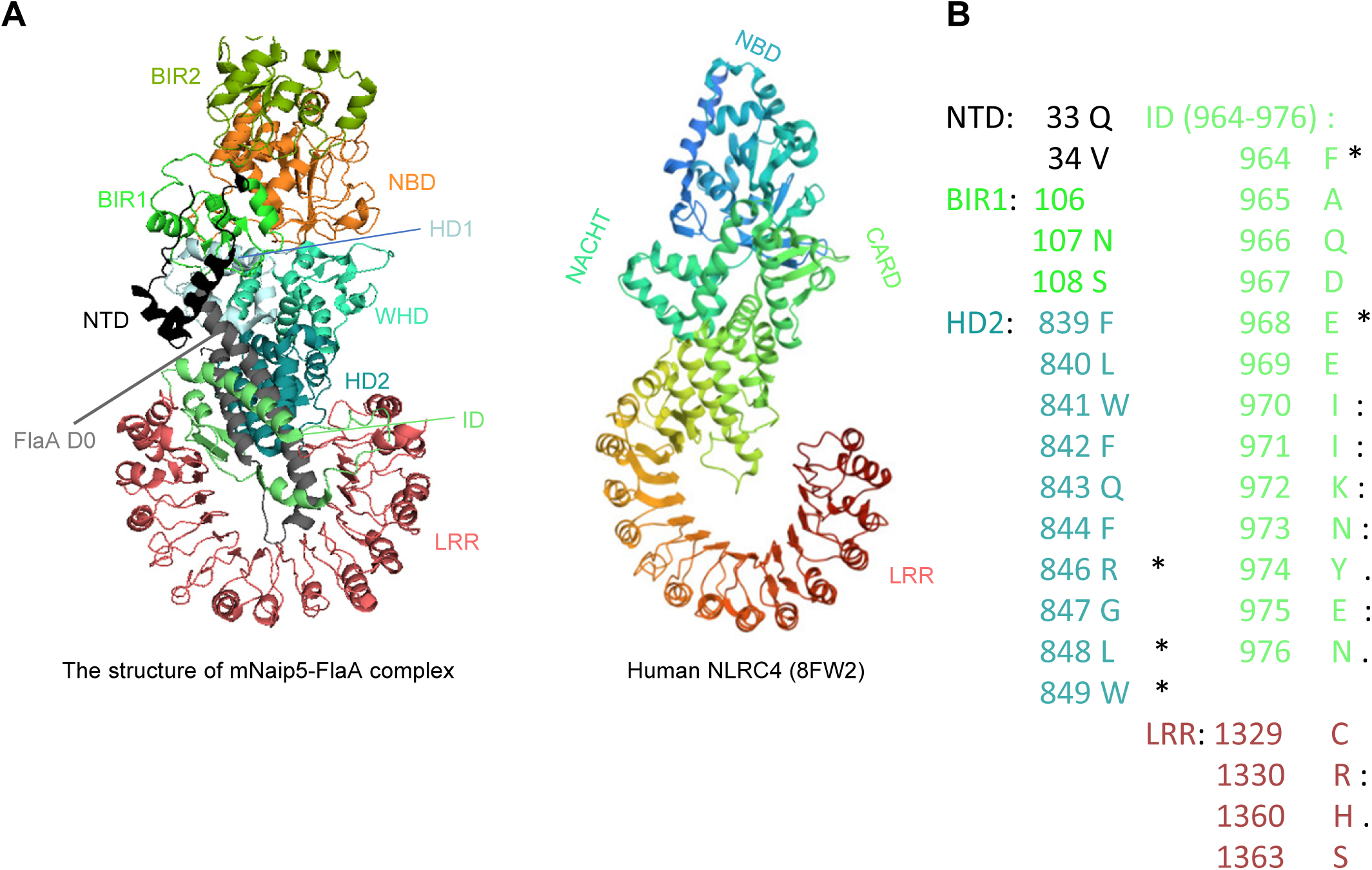
Structural and sequence similarities between mNaip5 and hNLRC4. **(A)** The published cryo-EM structures of the Naip5–flagellin complex (left, PDB: 5YUD) and human NLRC4 (right, PDB: 8FW2) are shown. **(B)** Critical residues in NAIP5 involved in flagellin recognition are distributed across different domains and are either conserved in, or share similar properties with, the corresponding residues in human NLRC4.

Additionally, a perspective comparing NAIP-NLRC4 sensing across species may offer a useful framework for understanding our findings. As mentioned above, unlike mice, humans and several other species possess only one NAIP. Thus, a single NAIP appears more common in nature. Although convergence of multiple functions in one species (e.g. mice) onto a single homologue in others is not unusual, the strict differences between mouse Naip ligand specificity would argue against a single hNAIP being able to cover *all* ligands sensed by multiple gene products in the murine system whilst retaining a necessary level of ligand specificity to prevent autoinflammation. The range of different reported T3SS components sensed by hNAIP – *Pseudomonas aeruginosa* PscI and PscF, *Salmonella* PrgI, and enterohemorrhagic *Escherichia coli* EprI ^(Yang, Zhao et al. 2013, Grandjean, Boucher et al. 2017, Matico, Yu et al. 2024)^ – would require hNAIP to accommodate a certain amount of structural flexibility only for T3SS components without losing specificity. Conceptually, it is therefore hard to envisage that the same hNAIP would not only be able to recognize different T3SS components but also specifically recognize a range of different flagellins, which mice employ to specific hNaips 1 and 2. In the absence of multiple human NAIPs, another receptor is thus plausible and our data suggest hNLRC4 has this capacity. One can only speculate why flagellin sensing at a genetic and functional level thus seems to be arranged differently in mouse vs man. One hypothesis could be that in humans hNLRC4 sensing provides redundancy for a single hNAIP sensor. Moreover, a preference to sense PAMPs over MAMPs might be at work: whereas multiple murine Naips feeding into a single murine Nlrc4 may treat the sensed MAMPs and PAMPs as relatively ‘equal’, the human system, by hNAIP being assigned to sense only a T3SS PAMP rather than the MAMP flagellin, may be genetically geared towards being more responsive to PAMP recognition. This is also consistent with the observation that in our hands, hNAIP, when present, seems to dampen flagellin sensing by NLRC4. We concede this potential regulatory function of hNAIP suggested by our data requires additional investigation but is noteworthy and would explain why multiple human cell types respond poorly to intracellular flagellin whereas their mouse counterparts do. Moreover, it would provide a functional enforcement of the preference for PAMP sensing over MAMP sensing. Under limited sensor resources, by prioritizing PAMP sensing via a single hNAIP whilst dampening the ‘noise’ of innocuous MAMP signals (Clasen, Bell et al. 2023), one could speculate this is the most reasonable and efficient strategy for humans to survive when faced with both bacterial pathogens and innocuous commensals.

## Materials

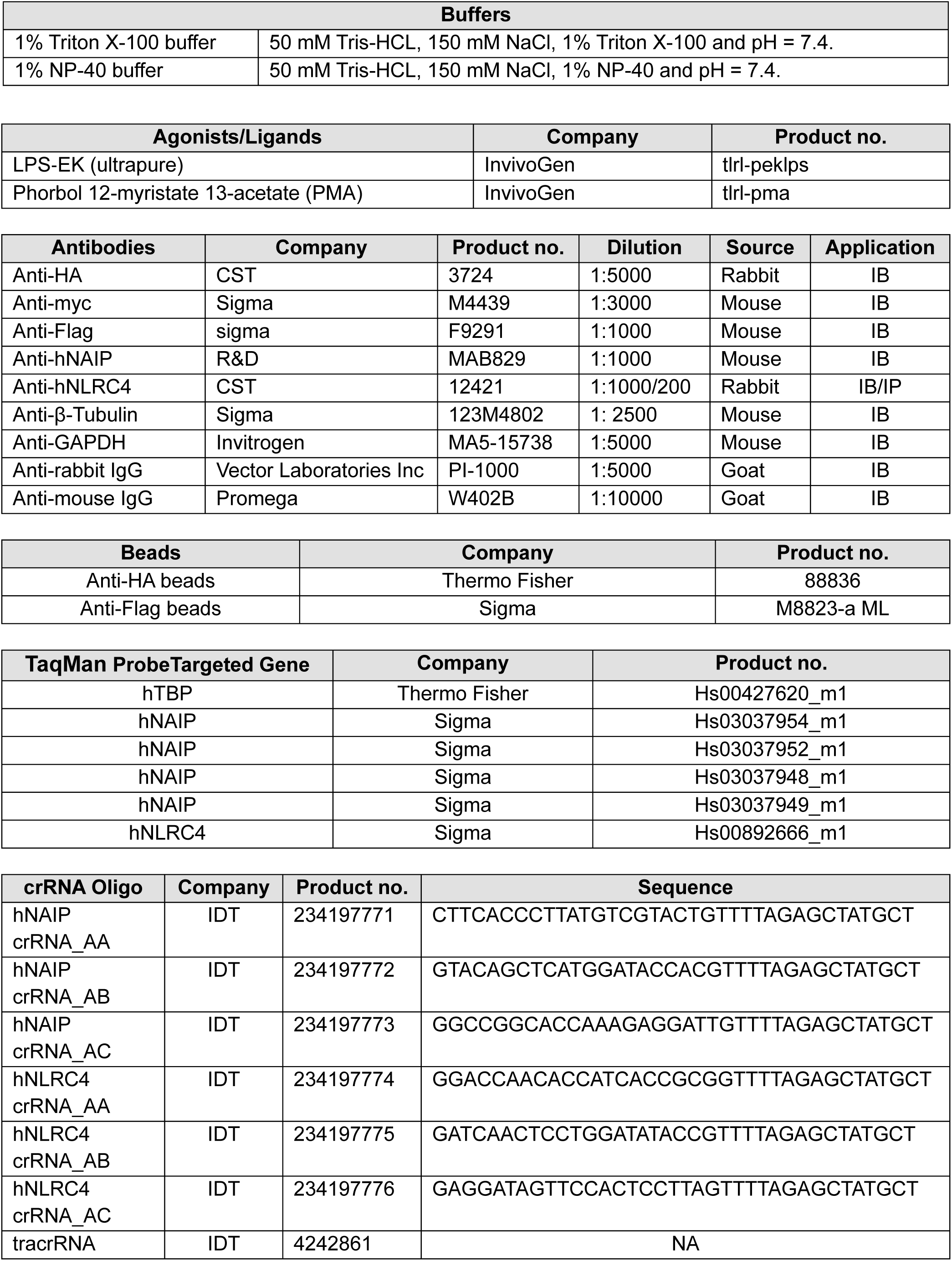

## Methods

### Cell culture

HEK293T cells were cultured in DMEM (Dulbecco’s Modified Eagle Medium)(Sigma, D5796-24X500ML) supplemented with 10% FBS (fetal bovine serum, Th. Geyer, 11682258), 2 mmol/L L-glutamine (Gibco, 2503008), 100 U/mL penicillin/streptomycin (Thermo Fisher, 15140122). HEK293T cells were grown at 37°C and 5% CO_2,_ and split 1:10 every 5 days or when confluence reached 80%. THP-1 WT and various knockout cell lines were cultured in RPMI-1640 medium (PAN Biotech, P04-18525). Supplemental reagents and culture conditions were the same as those used for HEK293T cells. The cells were split 1:5 every 5 days or when the density reached 10^6^ cells/mL. For stimulation, THP-1 cells were seeded into 24-well plates with 0.5 mL of pre-warmed RPMI-1640 per well and 100 ng/mL of PMA one night before. Next, the cell culture medium was replaced with fresh medium without PMA to rest the cells for 48 h. Then, the culture medium was changed again with fresh RPMI-1640 containing 100 ng/mL of LPS (Lipopolysaccharide) (InvivoGen, tlrl-peklps) for 3 h priming. In some conditions, the cells were incubated with 10 µM MCC950 (Selleckchem, S8930) for 1 h before LPS priming. And then the cell culture medium was removed and 0.5 mL of pre-warmed opti-MEM contained 10 µM of nigericin(InvivoGen, tlr-nig) was added per well for 1h. The supernatant was collected for further experiments.

### Plasmid isolation

Plasmid isolation was carried out using the Promega mini-prep (Promega, A1223) and midi-prep systems (Promega, A2492) according to the instructions from the manufacturer.

### Gateway cloning

Gateway cloning BP and LR reactions were performed using the kits from Thermo Fisher, Gateway™ BP Clonase™ II Enzyme mix (Thermo Fisher, 11789020) and Gateway™ LR Clonase™ II Enzyme mix (Thermo Fisher, 11791020).

### Site-directed mutagenesis

Site-directed mutagenesis was carried out using the QuikChange II XL Site-directed mutagenesis kit (Agilent Technologies, 200521) according to the manufacturer’s instruction. 1 µL of the Dpn I restriction enzyme (10 U/µL) was mixed with each PCR reaction, and the mixtures were pipetted up and down for several times. The tube was then incubated at 37°C for 1 h to digest the parental plasmid. The new plasmids were transformed into XL10-Gold ultracompetent cells (with kit) and mutations confirmed by Sanger sequencing.

### Enzyme linked immunosorbent assay (ELISA)

The human IL-1β ELISA kit (BioLegend, 437016) was from BioLegend, the human IL-18 ELISA kit (R&D, DY318-05) was from R&D and the human TNF ELISA kit was from Invitrogen (Invitrogen, 88-7346). ELISAs were performed based on the instructions from the manufacturers.

### SDS-PAGE electrophoresis

Typically, 50 µL of lysate was mixed with 20 µL of 4 x LDS Sample Buffer (Invitrogen, NP0008), 8 µL of 10 x Sample Reducing Agent (Invitrogen, NP0009) and 2 µL of ddH_2_O in a total volume of 80 µL. Samples were boiled at 95 °C for 5 min before loading 35 µL of protein sample per lane. Gels were run at 80 mV constant voltage for 30 min. Then, the voltage was increased to 120 mV for another 60 min.

### Immunoblot

Protein from SDS-PAGE gels were transferred to nitrocellulose membrane (Merck, GE10600003) using a Bio-Red Semi-Dry blotting system. Different protocols and time were used based on different molecular weights of the protein samples. After transfer, the membrane was blocked with 5% BSA (Bovine Serum Albumin Fraction V) (Biomol, 01400.250) or milk (Roth, T145.3) in Tris-buffered saline (TBS) with 0.1% Tween TBS-T wash buffer for 1 h at room temperature. Then, the primary antibodies were diluted with 5% BSA-containing wash buffer and the membranes incubated with the primary antibodies at 4 °C in a 50 mL Falcon while rotating overnight. Next, the membranes were washed three times for 5 min with 1x TBS-T. Washed membranes were incubated with secondary antibodies in 5% non-fat milk in TBS-T while rotating for 2 h at room temperature. Then, the membranes were washed with 1x TBS-T three times for 5 min. After washing, chemiluminescent substrate (Thermo Fisher, 34580) was added and the membranes imaged using an Odyssey FC imaging system (Licor). Blots were analyzed using Licor Imaging Studio Lite 5.2.

### Ready blue protein staining of gels

ReadyBlue™ Protein Gel Stain Solution (Sigma-Aldrich, RSB-1L) was purchased from Sigma-Aldrich and the following staining steps were also based on the instructions from manufacturer. Briefly, after electrophoresis, the gels were transferred from the cassette to a container and incubated with five times gel volume of Protein Gel Stain Solution with shaking at room temperature at least one hour. For higher sensitivity, the incubation time was prolonged to 2 h or even overnight. After staining, the gels were washed with ddH_2_O to remove any remaining solution. Gels imaging was done with a Vilber-Lourmat Fusion gel documentation system.

### RT-qPCR

RNA isolation was performed using an AllPrep DNA/RNA Mini Kit (Qiagen, 80234). 5 x 10^6^ –10^7^ cells were collected and centrifuged at 1600 rpm/min for 3 min to pellet cells and then supernatant was discarded. Then cells were lysed in 350 μL of buffer RLT (with kit) and incubated on ice for 2 min. Then lysate was centrifuged at 16100 x g for 3 min at 4 °C in a 2 mL Eppendorf tube. Next, the lysate was transferred to a clean 2 mL Eppendorf tube without disturbing the pellet. The following RNA isolation was done using the QIAcube robot (Qiagen) based on the instructions from the manufacturer. Genomic DNA contained in the isolated RNA samples was removed by using the digestion system from Thermo Fisher (AM1906). Reverse transcription was performed using a High-Capacity RNA-to-cDNA Kit (Thermo Fisher, 4387406). 9 µL of digested RNA was mixed with 10 µL of RT buffer and 1 µL of reverse transcriptase. The mixture was pipetted up and down several times. Meanwhile, a negative control was set up including 9 µL of digested RNA and 11 µL of ddH_2_O. The mixtures were incubated at 37 °C for 1 h and then heated up to 95°C for 5 min. Then the tube cooled down to 4°C. To perform the following qPCR experiment, the complementary DNA (cDNA) was diluted with 1:10 and 4.5 µL combined with 5 µL TaqMAN Universal MasterMix II (2x) (Thermo Fisher, 4440040), 0.5 µL TaqMAN Gene expression Assay (Probe and Primers, 20x). cDNA was replaced with ddH_2_O in the negative control group and primers for the house keeping gene TATA-box binding protein (*TBP*) were used as a reference gene. After pipetting, the plate was sealed and centrifuged briefly. Then the samples were run on a QuantStudio Real-Time PCR system. The relative quantification analysis was done by normalizing the target gene Ct value to the housekeeping gene *TBP* with the ΔΔCt method in Microsoft Excel 2019.

### Cytotoxicity detection

The cytotoxicity detection kit (Roche, 11644793001) was purchased, and the steps were performed based on the instructions from manufacturers. First, pure BSA standard samples were prepared and non-FBS containing samples were diluted properly. Then 100 µL of the standards or samples were loaded into 96-well plate. Mix LDH Reagent 1 and 2 (with kit) at ratio 1:45 based on the instructions. Then, 100 μL of the reagent mixture was loaded to each well and the 96-well plate was shaken briefly to mix well. The plate was incubated at room temperature for 5 min and meanwhile the color of the wells changed gradually. Then the LHD absorbance was detected under 490 nm by a standard plate reader.

### Purification of his-tagged Lfn-FlaA

To investigate the response of THP-1 to cytosolic flagellin, *Legionella pneumophila* flagellin FlaA was needed to purify at first. Therefore, his-tagged lethal factor N-terminal domain (Lfn)-FlaA expression plasmid in *E. coli* was generated based on ^(Rauch, Tenthorey^ ^et^ ^al. 2016)^ and then transformed into BL21 strain. FlaA expression was induced and then the bacteria were lysed. FlaA protein was captured by the anti-his tag column and purified with following size exclusion chromatography.

### Co-IP from transfected HEK293T cells

The indicated plasmids were transfected into HEK293T for 48 h. Then the culture medium was removed, and the plates were washed with cold 1x PBS. For each 10-cm dish, 850 µL of pre-chilled 1% Triton X-100 buffer containing protease and phosphatase inhibitors cocktail (Roche, 11836170001) was added to lyse the cells and the dish was placed on ice for 30 min to lyse completely. The plate was shaken several times during cell lysis. Lysate was then transferred from the plates to a pre-chilled 1.5 mL Eppendorf tube. The tube was centrifuged at 4°C with 16,000 x g for 15 min. Then the supernatant was collected and 50 µL of the supernatant was used to prepare the whole cell lysate (WCL) sample. 25 µL of the appropriate magnetic beads (e.g. anti-Flag or anti-HA) were pipetted to a new tube and the beads were washed with 0.4 mL of pre-chilled 1% Triton X-100buffer. The beads were then vortexed briefly and spun down. Then the tube was placed on the magnetic stand and left for 1 min to allow all the beads to adhere to the stand. Then the supernatant was removed and this washing step repeated one more time. Finally, the beads were resuspended with 25 µL of 1% Triton X-100 buffer and then the resuspended beads were added to the remaining lysate and the tube was rotated in the cold room overnight. On the second day, the tube was placed on the magnetic stand and left for 1 min. Then the lysate was removed. The beads were then washed with 1% Triton X-100 lysis buffer 3 times. After washing, the beads were resuspended in 50 µL of immunoblotting loading reagent and then boiled for 5 min at 95°C to detach the target protein from the beads.

### Co-IP in THP-1 cells

THP-1 cells were seeded into 10 cm dishes and cultured with 100 ng/mL PMA (phorbol 12-myristate 13-acetate) (InvivoGen, tlrl-pma) overnight to differentiate THP-1 cells towards adherent macrophage-like cells. Cells were lysed using pre-chilled 1% Triton X-100 buffer containing protease and phosphatase inhibitors (Roche, 11836170001) and centrifuged at 4°C with max speed for 15 min. Supernatant was collected and anti-hNLRC4 antibody (CST, 12421) was added to the lysate at a dilution of 1:200. The mixture was rotated in the cold overnight and 18 µL of protein G dynabeads (Invitrogen, 10003D) were pipetted into to each sample. The mixture was rotated in the cold room for another room. Then, the tube was placed on the magnetic stand for 1 min. Liquid was removed and then the beads were washed with 1% Triton X-100 lysis buffer 3 times. After washing, the beads were resuspended in 50 µL of immunoblotting loading reagent and then boiled for 5 min at 95 °C to detach the target protein from the beads.

### Native PAGE

HEK 293T cells were lysed in native lysis buffer, 50 mM Tris-HCL, 150 mM NaCl, 1% NP-40 and pH = 7.4. Lysates were then centrifuged at 16,000 x g for 30 min at 4 °C. Then the supernatant of the lysate was collected and mixed with the NativePAGE Sample buffer (Thermo Fisher, BN20032) following the instructions from manufacturer. The mixture was loaded into the pockets of 3–12% Bis-Tris native-PAGE gel (Thermo Fisher, BN1001BOX). The electrophoresis of NaitvePAGE was performed with NativePAGE running buffer (BN2001). After electrophoresis, the gel was incubated with 10% SDS buffer for 10 min at room temperature to denature and negatively charge proteins in the native-PAGE gel for transfer. The steps of protein transfer were the same as for conventional immunoblotting.

### CRISPR-Cas9-mediated generation of THP-1 knockout cell lines

Three clustered regularly interspaced short palindromic repeats (CRISPR) RNAs (crRNAs) were designed either for human NAIP or NLRC4 using the design tool available on the IDT company website. Cas9-mCherry expressing THP-1 cells were a kind gift from Dr. Hossam Gewaid, Trinity College Dublin. The crRNA (IDT, 234197771-6) and tracrRNA (IDT, 4242861) were resuspended in IDTE buffer (IDT, 11-01-02-02) respectively following the instructions from the manufacturer. 3.0 μL of crRNA and tracrRNA were mixed together and then heated at 95°C for 5 min. Then the mixture was cooled down to room temperature gradually. 4 μL of the oligo complex were then mixed with 6 μL of IDTE Buffer and kept on ice until used. crRNA-tracrRNA complex was delivered to the THP-1 cells by electroporation using a Cell Line Nucleofector™ Kit V (Lonza, VCA-1003). For each oligo complex, 82 µL of Solution Reagent and 18 µL of Supplement Reagent were mixed well by pipetting up and down gently. 10^6^ THP-1 cells were always collected and mixed with 100 µL of electroporation reagent as mentioned above. 10 µL of oligo complex was admixed to the cell suspension. Electroporation was performed using a Lonza Nucleofector 2b. The mixture of cell suspension with RNA oligos was transferred from the tubes to the cuvette and then the cuvette placed to the electroporation chamber in the nucleofector. The “X” button was pressed to start the program. After electroporation, the cell suspension was transferred to the prepared prewarmed 24-well plate containing 1 mL fresh RPMI-1640 and the cells then were incubated at 37°C, 5% CO_2_ for 5 days. Then the edited cell pools were transferred to a T25 bottle with 3 mL of culture medium to bulk up. When possible, aliquots of the pools were frozen. For generating single clones, the pools were transferred from T25 bottles to T75 bottles and incubated at 37°C, 5% CO_2_ for 1 week waiting for the cell concentration of these pools to reach 10^6^/mL. Then the pools were transferred into 15 mL Falcon tubes and centrifuged at 1,000 rpm/min for 5 min. The supernatant was removed, and the cell pellet was resuspended with 10 mL of fresh RPMI-1640. The cell number was counted and then resuspension was diluted to 10 cells per mL with a total volume of 55 mL. Several 96-well plates were labeled and 100 µL of diluted cell suspension was added to each well, which means each well contained one single edited cell on average. Then another 100 µL of fresh RPMI-1640 was added to each well. The growth of single clones was checked under the microscope every day and single clones could be easily found one month later. Then the single clones are identified by immunoblotting assay. Appropriate PCR primers annealing 5’ and 3’ of the gRNA sequence were used to perform PCR and the PCR products were purified and then Sanger sequenced. Sequencing results of knockout fragments were aligned with WT gene sequence using Geneious 6.0 and homozygous deletion or insertions leading to disruption of the ORF verified.

### Delivery of purified Lfn-FlaA via PA

PA was purchased from Alpha Diagnostic (Alpha Diagnostic, AV-9140-100) and Lfn-Needle was purchased from InvivoGen (Invivogen, tlrl-ndl), LFn-MxiH was the kind gift from Prof. Florian I. Schmidt, University of Bonn. Lfn and Lfn-FlaA were purified as mentioned above. 3×10^5^ THP-1 cells were seeded and differentiated as mentioned above. At second day, reagents were diluted and pipetted with prewarmed Opti-MEM to the desired working concentration. PA is 1 µg/mL, Lfn, Lfn-MxiH and Lfn-Needle are 0.1 µg/mL, Lfn-FlaA is 0.2 µg/mL. Then the reagents were added into the culture supernatant and incubated for the desired time. The culture medium was collected for the detection of the concentrations of human IL-1β and IL-18.

### Bacterial infection of THP-1 cells

*Salmonella Typhimurium* (from Prof. Samuel Wagner, University of Tübingen) bacteria strain was streaked out, and single clones were picked from the plate and cultured in LB medium with 50 µg/mL of streptomycin overnight. Bacterial culture was diluted to OD_600_ = 0.2 and then incubated with shaking for 2 h to induce the expression of SPI-1 (Salmonella pathogenicity island 1). THP-1 cells were seeded and differentiated into 24-well plate as mentioned above. Then cells were primed with 50 ng/mL of LPS for 3 h. Prior to infection, the bacteria were diluted to achieve an MOI of 10. Then, bacteria were added to each well and then the plate was centrifugated with 500 g for 10 min under 22°C. Then the plate was incubated at 37 °C for another 30 min. Infected THP-1 cells were washed three times with PBS and fresh media containing 100 ng/ml gentamycin was added to each well to kill the extracellular bacteria. At the desired timepoint, cell culture medium was harvested and subjected to ELISA assays. *Legionella pneumophila* (provided by Prof. Samuel Wagner, University of Tübingen but originally obtained from Prof. Russel Vance, University of California, Berkeley) infection assay was performed with the similar steps as mentioned above.

### Reconstitution of the human NLRC4 inflammasome in HEK293T

In brief, HEK 293T cells were seeded in 6-well plates with 1 mL DMEM one night before. On the second day, different expression constructs were transfected using lipofectamine 2000: 100 ng of FlaA, 100 ng of hNAIP, 200 ng of hNLRC4, 100 ng of pro-hCaspase1 and 200 ng of pro-hIL1β. After 6 h of incubation, the transfection reagents were removed and 2 mL of fresh DMEM was added per well. After 48 h of incubation, the culture medium was collected for detection of cytokine release by ELISA. The cells were washed with pre-cold 1x PBS once and lysed with 300 μL of 1% NP-40 buffer. The lysate was centrifuged at 16100 g for 30min with 4 °C and then the supernatant was collected for native PAGE assay or normal immunoblotting analysis.

### Immunofluorescence (IF) staining

HEK293T cells were seeded and transfected with the NLRC4 inflammasome components as shown in the figures. Especially, FlaA was labeled with mCherry and hNLRC4 was labeled with eGFP. Then cells were fixed with Fixation Buffer (BioLegend, 420801) for 15 min at room temperature. Then the coverslips were washed twice with 1 x PBS by gently shaking the plate, and 5 min for each washing step. Hoechst 33342 (Thermo Fisher, H21492) was diluted in 1:10000 with 1 x PBS and then the coverslips were incubated with diluted Hoechst for 10 min at room temperature in the dark. Then the coverslips were washed with 1 x PBS one time. The coverslips were dipped in distilled water to remove salt and then placed on a soft tissue paper to dry briefly. The coverslips were mounted in 10 µL of mounting solution (Thermo Fisher, P36961). Then coverslips were gently lowered down and incubated at room temperature on a straight surface for drying overnight. Then they were kept at 4°C until microscopy was performed and for longer-term storing.

## Data analysis and statistics

Analysis of DNA sequencing results and the editing of plasmid maps was done with Geneious 6.0. DNA agarose gels imaging was performed with Fusion SL software (Vilber Lourmat). Analysis of immunoblotting was done using Image Studio (Li-COR). Microcopy data was analysis with ImageJ. qPCR results were analyzed by QuantStudio Real-Time-PCR software version 13 and Excel (Microsoft, 2019). Data were tested for normality first and then *p* values were calculated using Excel and GraphPad Prism version 8.0 with Student’s *t*-tests or one-way ANOVA as indicated in figure legends.

## Acknowledgements

We thank Maria Mateo Tortola for general microscopy support, Michael Braun (Stehle/Hartmann lab, University of Tübingen) for assisting with flagellin protein purification, Samuel Wagner (Institute of Medical Microbiology, University of Tübingen) for preparing *Salmonella* Typhimurium and *Legionella pneumophila* strains which were a kind gift of Russel Vance (University of California, Berkeley, USA). We gratefully acknowledge receipt of Lfn plasmids from Isabella Rauch as described in (Rauch, Tenthorey et al. 2016). We also thank Hossam Gewaid (Trinity College Dublin, Ireland) for kindly sharing of Cas9-mCherry expressing THP-1 cells.

## Funding

The study was supported by the funds from the Volkswagenstiftung Momentum grant “InnatelyHuman” (Project 0070643-00; to X.L. and A.N.R.W.), DFG (Deutsche Forschungsgemeinschaft/German Research Foundation) via the Clusters of Excellence “iFIT – Image Guided and Functionally Instructed Tumor Therapies” (EXC-2180, to A.N.R.W) and “CMFI – Controlling Microbes to Fight Infections” (EXC-2124, to G. L. and A. N. R. W.) and via grant We-4195/18-1 (to A.N.R.W.). We also gratefully acknowledge support by the University of Tübingen Teach@Tübingen program, the University Library Tübingen and the Medical Faculty Library open access funds.

## Author contributions

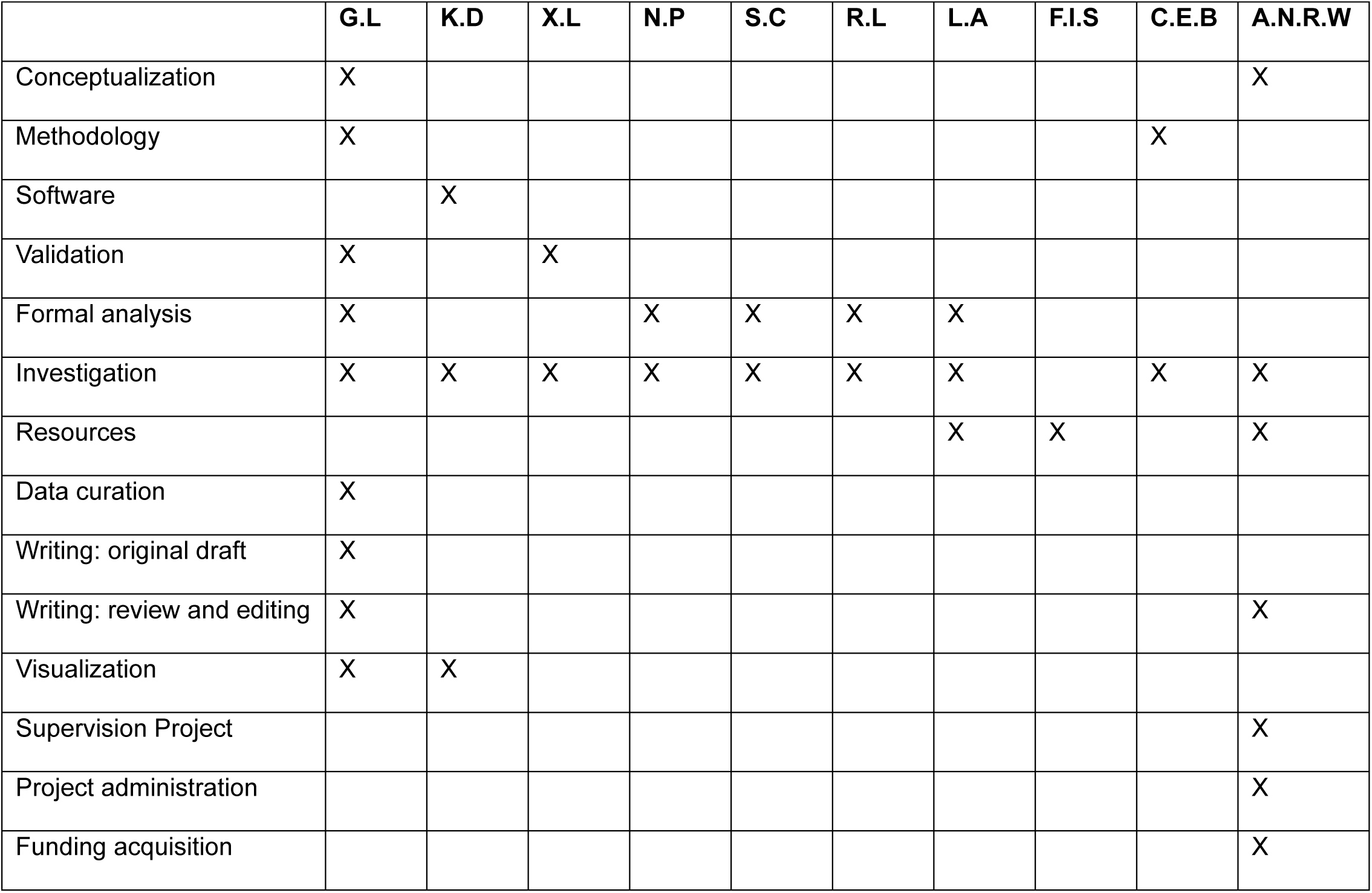

## Figure Legends

**Figure S1: mRNA and protein levels of the hNAIP and hNLRC4 in HEK293T and THP-1 cells. (A)** Schematic of the alignment between LhNAIP and ShNAIP. **(B)** Different TaqMan qPCR primers, specifically targeting the hNAIP exon 13 DNA sequence, were used to measure the mRNA levels of LhNAIP, labeled as LhNAIP-1 and LhNAIP-2. Another TaqMan primer, specifically targeting the beginning of the hNAIP gene, which is shared between LhNAIP and ShNAIP, was used to detect the total hNAIP mRNA levels, and hence labeled as Total hNAIP. The mRNA levels of hNAIP were normalized to the mRNA levels of the housekeeping gene, TBP. **(C)** The relative mRNA levels of hNLRC4 in HEK293T and PMA-differentiated macrophage-like THP-1 cells are shown. **(D)** Immunoblot analysis of protein levels of total hNAIP and hNLRC4 in HEK293T and THP-1. Data are representative of at least n=2 independent biological replicates.

**Figure S2: Flagellin-mediated IL-β release in the reconstituted hNLRC4 inflammasome system. (A-B)** hNLRC4 inflammasome reconstitution in HEK293T cells. HA-flagellins, Flag-ShNAIP, myc-hNLRC4, pro-hCaspase-1 and pro-hIL-1β expression plasmids were co-transfected as indicated. After 48 h, hIL-1β release was measured by triplicate ELISA. Data are representative of n = 3 independent biological replicates (mean+/-SD shown; each dot represents one technical replicate; p-values according to Student’s *t*-test).

**Figure S3: Detection of hNAIP and hNLRC4 in the knockout cell lines. (A-B)** IB of the protein levels hNAIP and hNLRC4 both in WT and different knockout cell pool WCLs. **(C-D)** IB of the protein levels hNAIP and hNLRC4 both in WCLs from WT and different single knockout clones picked up from the corresponding pool cell lines. In **(A)** hNAIP KO pool cell line generated with gRNA AA (Lane 2) and in **(B)** hNLRC4 KO pool cell line generated with gRNA AC (Lane 4) were used in Figure 4B with two more independent repeats. Knockout efficiency with single clones in **(C)** and **(D)** was verified with one independent experiment.

**Figure S4: hNLRC4 interacts with and is inhibited by hNAIP. (A)** hNLRC4 inflammasome reconstitution in HEK293T cells as in Fig. 4G but with an increased concentration of an GFP construct instead of hNAIP. **(B)** Lysates from A analyzed using SDS-PAGE and IB using the indicated antibodies. Data are representative of n = 3 independent biological replicates (mean+SD; each dot represents one technical replicate, Student’s *t*-test).

**Figure S5: The activation of the hNLRC4 inflammasome pathway in WT and knockout cell lines induced by Needle and MxiH. (A-C)** WT and KO PMA-differentiated THP-1 cells lines were stimulated with PA alone or combined with Lfn, LFn-MxiH, and Lfn-Needle. 2 h later, released cytokines and LDH were quantified in technical triplicates. Data are representative of n=3 independent biological replicates (mean+SD, each dot represented one technical repeat).

**Figure S6: Protein sequence alignment between human NAIP, hNLRC4 and murine Naips. (A)** A phylogenetic tree was generated by Clustal Omega using the protein sequence alignment of hNAIP and murine Naip family. The “Neighbor Joining” method and uncorrected “P” were used to calculate the pairwise value. (B) Sequence alignment of the hNLRC4) and mNaip5 amino acid sequences generated using CLUSTAL O (v1.2.4).

## Notes

### Competing Interest Statement

The authors have declared no competing interest.

### Summary of Updates

Corrected a mistake in the labeling of Fig. 2B and 2C

