## Supplementary figures S1-S6 for "Human NLRC4 can act as a direct sensor for cytosolic flagellin"

### Figure S1

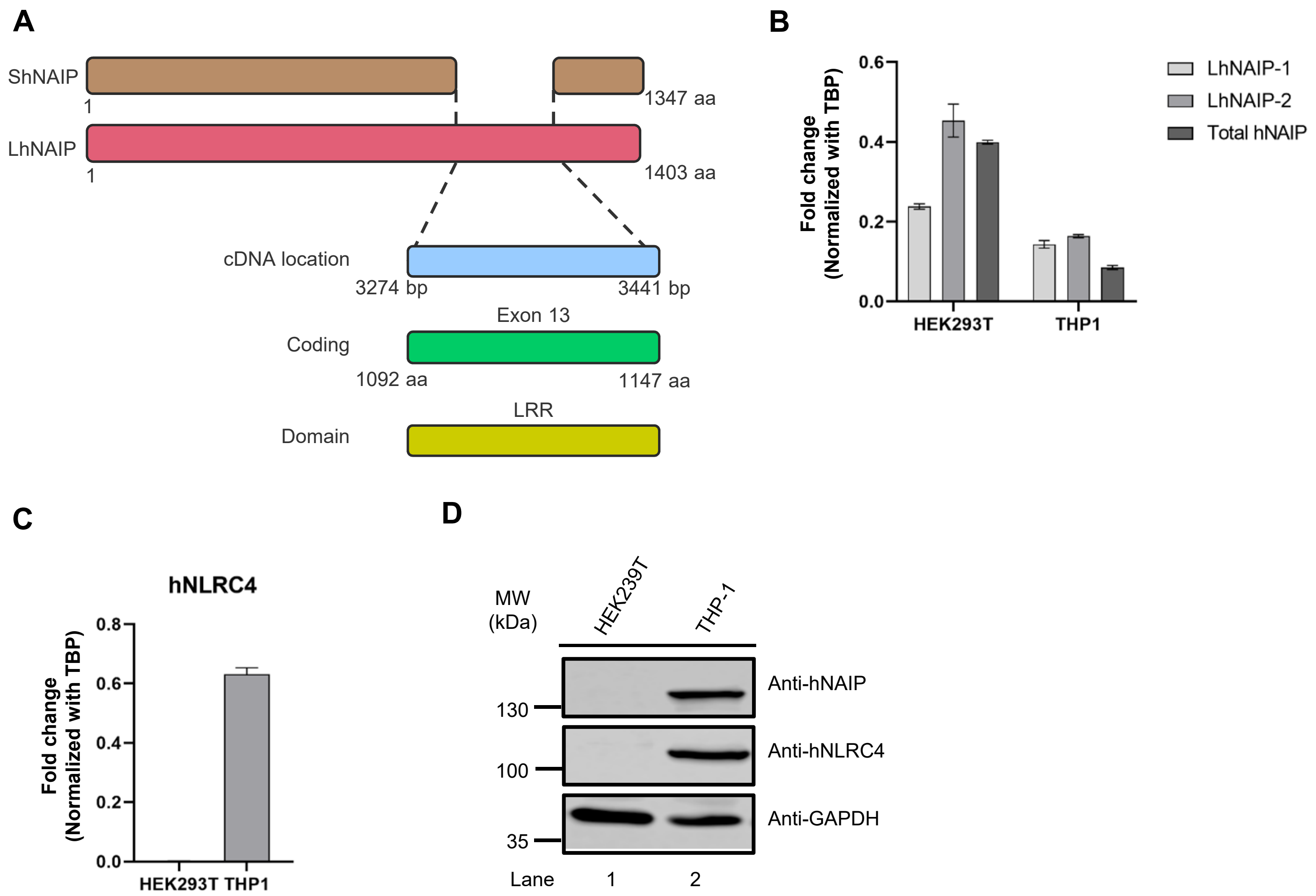
A
ShNAIP
1
1347 aa
LhNAIP
1
1403 aa
cDNA location
3274 bp
3441 bp
Exon 13
Coding
1092 aa
1147 aa
LRR
Domain
B
Fold change (Normalized with TBP)
0.6
0.4
0.2
0.0
HEK293T
THP1
LhNAIP-1
LhNAIP-2
Total hNAIP
C
hNLRC4
Fold change (Normalized with TBP)
0.8
0.6
0.4
0.2
0.0
HEK293T
THP1
D
MW (kDa)
HEK239T
THP-1
Anti-hNAIP
Anti-hNLRC4
Anti-GAPDH
130
100
35
Lane
1
2

#### Figure S2

**A**

| Group | 1 | 2 | 3 | 4 |
|---|---|---|---|---|
| Flagellin | – | + | – | + |
| ShNAIP | – | – | + | + |
| hNLRC4 | + | + | + | + |
| pro-hCaspase1 | + | + | + | + |
| pro-hIL1β | + | + | + | + |

**B**

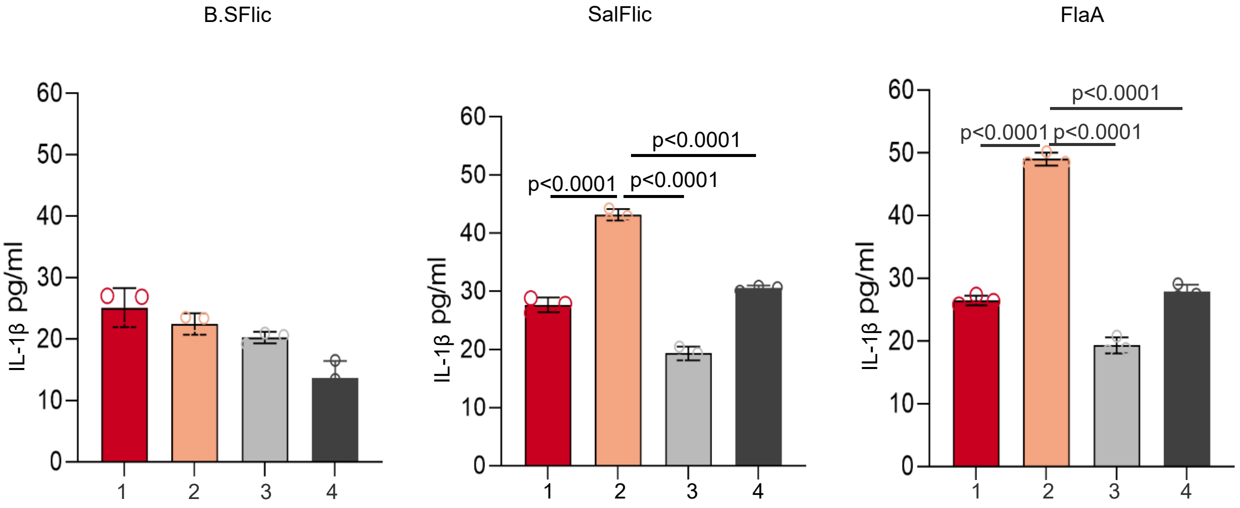

### Figure S3

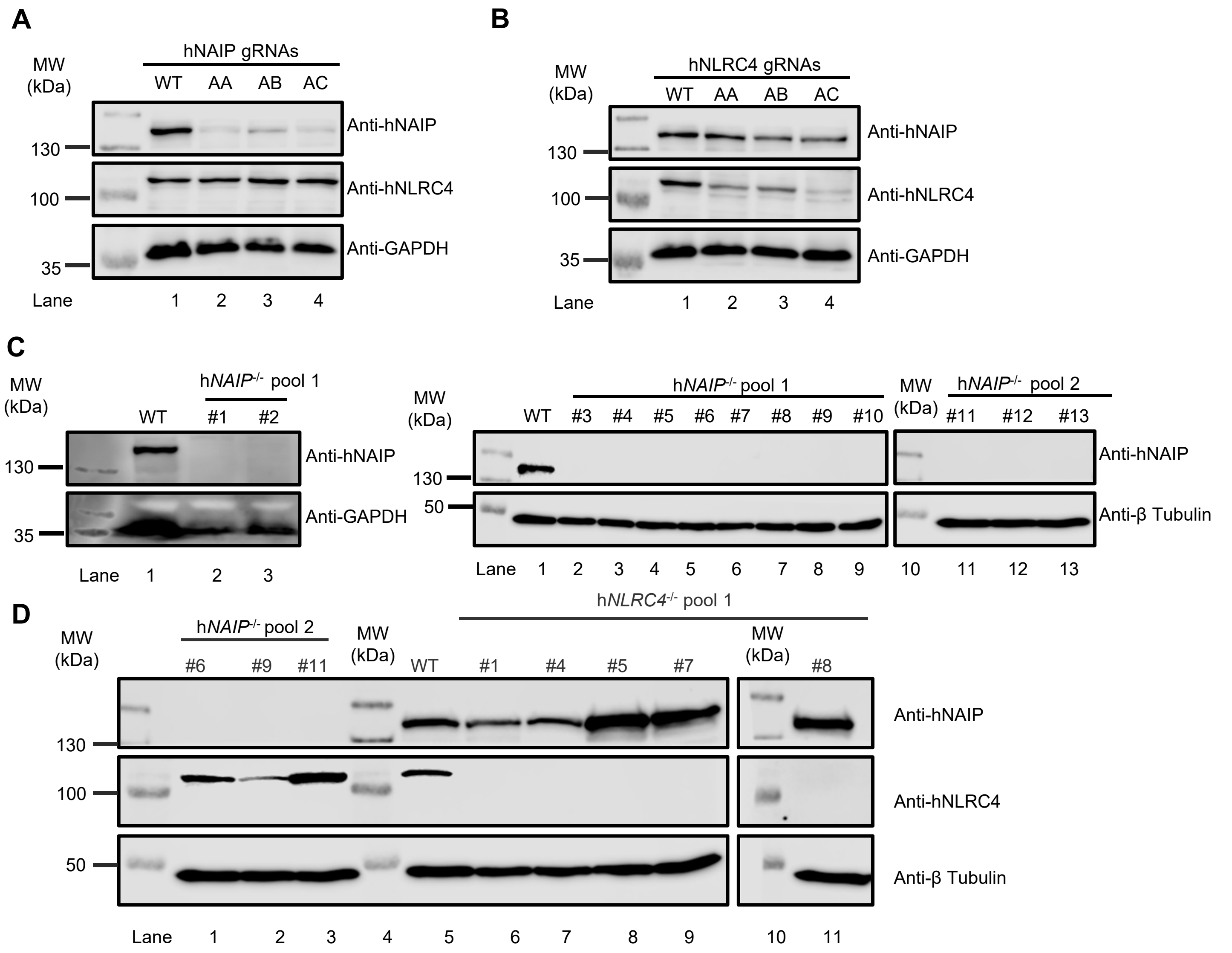
A
MW (kDa)
hNAIP gRNAs
WT AA AB AC
Anti-hNAIP
Anti-hNLRC4
Anti-GAPDH
130
100
35
Lane 1 2 3 4
B
MW (kDa)
hNLRC4 gRNAs
WT AA AB AC
Anti-hNAIP
Anti-hNLRC4
Anti-GAPDH
130
100
35
Lane 1 2 3 4
C
MW (kDa)
hNAIP-/- pool 1
WT #1 #2
Anti-hNAIP
Anti-GAPDH
130
35
Lane 1 2 3
hNAIP-/- pool 1
WT #3 #4 #5 #6 #7 #8 #9 #10
hNAIP-/- pool 2
#11 #12 #13
Anti-hNAIP
Anti-β Tubulin
130
50
Lane 1 2 3 4 5 6 7 8 9 10 11 12 13
D
MW (kDa)
hNAIP-/- pool 2
#6 #9 #11
hNLRC4-/- pool 1
WT #1 #4 #5 #7 #8
Anti-hNAIP
Anti-hNLRC4
Anti-β Tubulin
130
100
50
Lane 1 2 3 4 5 6 7 8 9 10 11

#### Figure S4

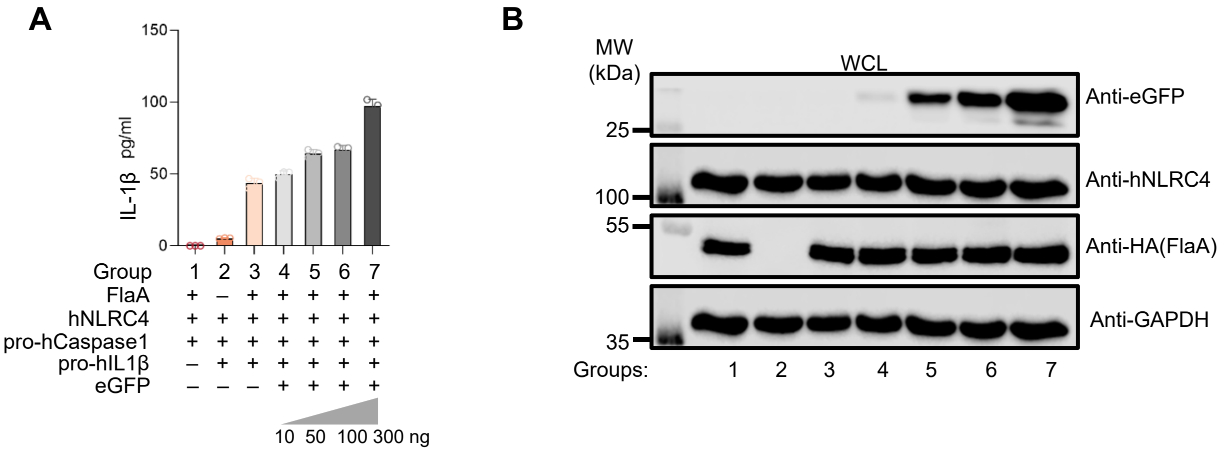
A
150
100
50
0
IL-1β pg/ml
Group 1 2 3 4 5 6 7
FlaA + – + + + + +
hNLRC4 + + + + + + +
pro-hCaspase1 + + + + + + +
pro-hIL1β – + + + + + +
eGFP – – – + + + +
10 50 100 300 ng
B
MW (kDa)
WCL
25
Anti-eGFP
100
Anti-hNLRC4
55
Anti-HA(FlaA)
35
Anti-GAPDH
Groups: 1 2 3 4 5 6 7

### Figure S5

**A**

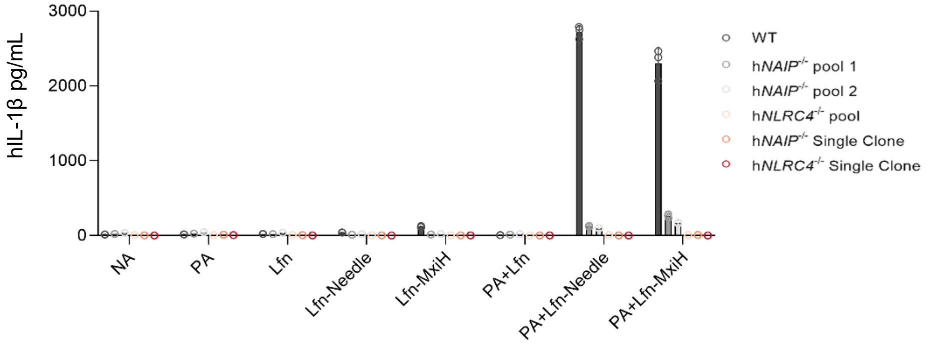
hIL-1β pg/mL
3000
2000
1000
0
NA
PA
Lfn
Lfn-Needle
Lfn-MxiH
PA+Lfn
PA+Lfn-Needle
PA+Lfn-MxiH
WT
hNAIP-/- pool 1
hNAIP-/- pool 2
hNLRC4-/- pool
hNAIP-/- Single Clone
hNLRC4-/- Single Clone

**B**

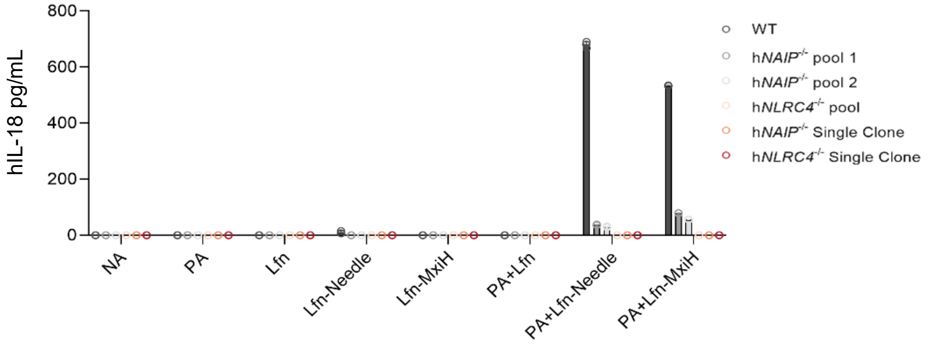
hIL-18 pg/mL
800
600
400
200
0
NA
PA
Lfn
Lfn-Needle
Lfn-MxiH
PA+Lfn
PA+Lfn-Needle
PA+Lfn-MxiH
WT
hNAIP-/- pool 1
hNAIP-/- pool 2
hNLRC4-/- pool
hNAIP-/- Single Clone
hNLRC4-/- Single Clone

**Figure S6**

**A**

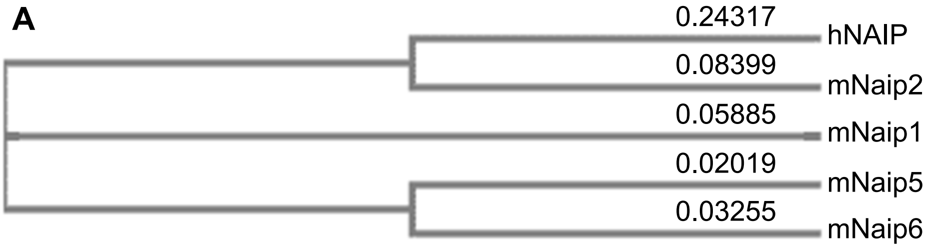


**B**

```
hNLRC4        ------------------------------------------------------------     0
mNaip5        MAEHGESSEDRISEIDYEFLPELSALLGVDAFQVAKSQEEEEHKERMKMKKGFNSQMRSE    60

hNLRC4        ------------------------------------------------------------     0
mNaip5        AKRLKTFETYDTFRSWTPQEMAAAGFYHTGVRLGVQCFCCSLILFGNSLRKLPIERHKKL   120

hNLRC4        ------------------------------------------------------------     0
mNaip5        RPECEFLQGKDVGNIGKYDIRVKRPEKMLRGGKARYHEEEARLESFEDWPFYAHGTSPRV   180

hNLRC4        ------------------------------------------------------------     0
mNaip5        LSAAGFVFTGKRDTVQCFSCGGSLGNWEEGDDPWKEHAKWFPKCEFLQSKKSSEEIAQYI   240

hNLRC4        -----------MNFIKDNSRAL--------------IQRMGMTVIKQ-----------IT    24
mNaip5        QSYEGFVHVTGEHFVKSWVRRELPMVSAYCNDSVFANEELRMDMFKDWPQESPVGVEALV   300
                          :*:*.  *                 :.: * ::*:           :.

hNLRC4        DDLFVWNVLNREEVNIICCEKVEQDAARGIIHMILKKG-SESCNLFLKSLK---------    74
mNaip5        RAGFFYT-GKKDIVRCFSCGGCLEKWAEGDDPMEDHIKFFPEC-VFLQTLKSSAEVIPTL   358
                 *.:.  ::: *. :.*    :. *.*   *  :     .* :**::**

hNLRC4        --EWNYPLFQDLNGQSLFHQTSEGDL---------------------DDLAQDLKDLYHT   111
mNaip5        QSQYALPEATET-----TRESNHGDAAAVHSTVVDLGRSEAQWFQEARSLSEQLRDNYTK   413
                ::  *   :       :::..**                       .*:::*:* * .

hNLRC4        PSFLNFY------PLGEDIDIIFNLKSTFTEPVLWRKDQHHHRVEQLTLNGLLQALQSPC   165
mNaip5        ATFRHMNLPEVCSSLGTDHLLSC-------DVSIISKHISQPVQEALTIPEVFSNLNSVM   466
               :* ::        ** *  :         :  :  *.  :   * **:  ::. *:*

hNLRC4        IIEGESGKGKSTLLQRIAMLWGSGKCKALTKFKFVFFLRLSR--AQGGLFETLCDQLLDI   223
mNaip5        CVEGETGSGKTTFLKRIAFLWASGCCPLLYRFQLVFYLSLSSITPDQGLANIICAQLLGA   526
               :***:*.**:*:*:***:**.** *  * :*::**:* **    : ** : :* ***.

hNLRC4        PGTIRKQTFMAMLLKLRQRVLFLLDGYNEFKPQNCPEIEALIKENHRFKNMVIVTTTTEC   283
mNaip5        GGCISEVCLSSSIQQLQHQVLFLLDDYSGLASLPQA-LHTLITKNYLSRTCLLIAVHTNR   585
               * * :  : : : :*:::******.*. :       :.:**.:*:  :. ::::. *:

hNLRC4        LRHIRQFGALTAEVGDMTEDSAQALIREVLIKE--LAEGLLLQIQKSRCLRNLMKTPLFV   341
mNaip5        VRDIRLYLGTSLEIQEFPFYNTVSVLRKFFSHDIICVEKLIIYFIDNKDLQGVYKTPLFV   645
              :*.** : . : *: ::   .: :::*:.: ::   .* *:: :  ..:*:.: ******

hNLRC4        VITCAIQMGESEFHS-HTQTTLFHTFYDLLIQKNKHKHKGVAASDFIRSLDHCGDLALEG   400
mNaip5        AAVCTDWIQNASAQDKFQDVTLFQSYMQYLSLKYK-----ATAEPLQATVSSCGQLALTG   700
              . .*:  : ::. :. . :.***::: : *  * *     .:*. :  ::. **:*** *

hNLRC4        VFSHKFDFELQDVS--SVNEDVLLTTGLLCKYTAQRFKPKYKFFHKSFQEYTAGRRLSSL   458
mNaip5        LFSSCFEFNSDDLAEAGVDEDEKLTTLLMSKFTAQRLRPVYRFLGPLFQEFLAAVRLTEL   760
              :**  *:*: :*::  .*:**  *** *:.*:****::* *:*:   ***: *. **:.*

hNLRC4        LTSHEPEEVTKGNGYLQKMVSISDITSTYSSLLRYTCGSS-VEATRAVMKHLAAVYQHG-   516
mNaip5        LSSDRQEDQDLGLYYLRQIDSPLKAINSFNIFLYYVSSHSSSKAAPTVVSHLLQLVDEKE   820
              *:*.. *:   *  **::: *  .  .::. :* *... *  :*: :*:.**  : :.

hNLRC4        CLLGLS----------------IAKRPLWRQESLQSVKNTTE----QEILKAININSFVE   556
mNaip5        SLENMSENEDYMKLHPQTFLWFQFVRGLWLVSPESSSSFVSEHLLRLALIFAYESNTVAE   880
              .* .:*                   * **  .  .* . .:*      :: * : *:..*

hNLRC4        CGIHLYQESTSKSALSQEFEAFFQGKSLYINSGNIPDYLFDFFEHLPNCASALDFIKLDF   616
mNaip5        CSPF--------------ILQFLRGKTLALRVLN-----LQYFRDHPESLLLLRSLKVSI   921
              *. .              :  *::**:* :.  *     :::*.. *:.   *  :*:.:

hNLRC4        YGGAMASWEKAAEDTGGIHME--EAPETYIPSRAVSLFFNWKQEFRTLEVTLRDFSKLNK   674
mNaip5        NGNKMSSYVDYSFKTYFENLQPPAIDEEY--TSAFEHISEWRRNFAQDEEIIKNYENIRP   979
               *. *:*: . : .*   :::     * *  : *.. : :*:::*   *  ::::.::.

hNLRC4        Q-----------------------------------DIRYLGKIFSSATSLRLQIKRCAG   699
mNaip5        RALPDISEGYWKLSPKPCKIPKLEVQVNNTDAADQALLQVLMEVFSASQSIEFRLFNSSG  1039
              :                                    :: * ::**:: *:.::: ..:*

hNLRC4        VAGSLSLVLSTCKNIYS-LMVEASPLTIEDERHITSVTNLKTLSIHDLQNQRLPGGLTDS   758
mNaip5        FLESICPALELSKASVTKCSMSRLELSRAEQELLLTLPALQSLEV--SETNQLPEQLFHN  1097
              .  *:. .*. .*   :   :.   *:  ::. : ::  *::*.:   :.::**  * ..

hNLRC4        LGNLKNLTKL----------------------IMDNIKM---NEEDAIKLAEGLKNLKKM   793
mNaip5        LHKFLGLKELCVRLDGKPNVLSVLPREFPNLLHMEKLSIQTSTESDLSKLVKFIQNFPNL  1157
              * :: .*.:*                       *:::.:   .*.*  **.: ::*: ::

hNLRC4        CLFHLTHLSDIGEGMDYIVKSLSSEPCDLEEIQLVSCCLSANAVKILAQNLHNLVKLSIL   853
mNaip5        HVFHLKCD--FLSNCESLMA-VLASCKKLREIEFSGRCFEAMTF---VNILPNFVSLKIL  1211
               :***.    : ..  : ::  : :.  .*.**:: . *:.* :.   .: * *:*.*.**

hNLRC4        DLSENYL-EKDGNEALHELIDRMNVLEQLTALMLPWGCDVQGSLSSLLKHLEEVPQLVKL   912
mNaip5        NLKDQQFPDKETSE---KFAQALGSLRNLEELLVPTGDGIHQVAKLIVRQCLQLPCLRVL  1268
              :*.:: : :*: .*   :: : :. *.:*  *::* * .::   . ::::  ::* *  *

hNLRC4        GLKNWRLTDTEIRILGAFFGKNPLKNFQQLNL-AGNRVSSDGWLAFMGVFENLKQLVFFD   971
mNaip5        TFHDILDDDSVIEIARAA-TSGGFQKLENLDISMNHKITEEGYRNFFQALDNLPNLQELN  1327
               :::    *: *.*  *   .. ::::::*::  .::::.:*:  *: .::** :*  ::

hNLRC4        FS---TKEFLPDPALVRKLSQVLSKLTFLQEARLVGWQFDDDDLSVITGAFKLVTA----  1024
mNaip5        ICRNIPGRIQVQATTVKALGQCVSRLPSLIRLHMLSWLLDEEDMKVINDVKERHPQSKRL  1387
              :.     .:  : : *: *.* :*:*  * . :::.* :*::*:.**... :

hNLRC4        ----------------   1024
mNaip5        IIFWKLIVPFSPVILE   1403
```
